# A selective top-down pathway from anterior cingulate cortex embeds a representation of saliency in hippocampus

**DOI:** 10.1101/2025.06.26.661852

**Authors:** Yaolong Li, Kotaro Mizuta, Steven J. Middleton, Yasunori Hayashi

**Affiliations:** Department of Pharmacology, Kyoto University Graduate School of Medicine, Kyoto 606-8501, Japan; Department of Biology, Division of Science, New York University Abu Dhabi, Abu Dhabi, United Arab Emirates

**Keywords:** anterior cingulate cortex, retrosplenial cortex, reward, place cells, hippocampus

## Abstract

For maximizing survival, animals must accurately memorize the structure of their environment. This is achieved by stabilizing and enriching hippocampal place cells near salient features, while the representation of neutral locations incrementally drifts with time. However, the circuit mechanisms by which top-down and bottom-up inputs selectively embed saliency into the hippocampal spatial map remain poorly understood. Here we identified a specific top-down input from the anterior cingulate cortex (ACC) to the dysgranular zone of the retrosplenial cortex (RSCd) that becomes active exclusively at reward locations. Activity of these ACC neurons is both necessary and sufficient for shaping the hippocampal saliency map. Unlike dopaminergic neurons in the ventral tegmental area, which respond to reward irrespective of the context, this class of ACC neurons encodes reward-context associations. Their optogenetic activation induces a reduction in locomotor speed, a behavioral correlate of saliency detection, while their inhibition disrupts the formation of the hippocampal place map. This work not only provides the first mechanistic insight into how context-dependent saliency is integrated in a top-down fashion into the hippocampal map to guide adaptive behavior but also challenges the classical view of memory consolidation theory where memory unidirectionally moves in bottom-up fashion from the hippocampus to the neocortex.

## INTRODUCTION

The ability to form a member on environmental saliency such as locations of food, threats, and other landmarks is crucial for maximizing an animal’s survival by enabling effective foraging, interactions with conspecifics and threat avoidance (Corkin, 2002; Goshen et al., 2011; Olton et al., 1979; Uekita, 2011). In the brain, external space is represented by place cells that activate when an animal passes through specific locations, a phenomenon that has been extensively studied in the hippocampus (Moser, 2015; O’Keefe, 1987). Saliency in an environment can influence the distribution of place cells within the hippocampal map of space, leading to an overrepresentation of the locations with enhanced saliency (Gauthier et al., 2018; Kaufman et al., 2020; Sato et al., 2020; Schuette et al., 2020). Importantly, this process is impaired in animal models of neurocognitive and neurodevelopmental disorders (Sato et al., 2020; Takamura et al., 2021).

The enrichment of representations around salient locations is in part accomplished by a selective stabilization of place cells near salient locations across days, while neutral locations gradually remap over time (Gauthier et al., 2018; Sato et al., 2020). However, how such overrepresentation is established has not been fully elucidated. One mechanism involves subcortical neuromodulatory inputs such as serotonergic inputs from medial raphe and noradrenergic and dopaminergic inputs from the locus coeruleus (Kaufman et al., 2020; Luchetti et al., 2020). Studies have shown that these neuromodulatory inputs to the hippocampus modulate reward-induced behavior and contribute to the overrepresentation of place cells near salient locations (Kaufman et al., 2020; Kempadoo, et al., 2016; Luchetti et al., 2020; Takeuchi et al., 2016).

In contrast, the role of top-down signaling from associative cortices to the hippocampus in saliency representation remains poorly understood. Among the various cortical regions sending projections to the hippocampus, the anterior cingulate cortex (ACC) is of particular interest as it is central to memory consolidation (Frankland et al., 2005; Rajasethupathy et al., 2015) and a top-down signal from ACC to hippocampus has been implicated in memory recall (Bian et al., 2019; Rajasethupathy et al., 2015; Yadav et al., 2022). Importantly, the ACC encodes a wide range of internal and external information including key components of saliency such as emotion, decision, and sensory features (Akam et al., 2021; Bota et al., 2021; Bush et al., 2002; Chien, et al., 2023; Hyman et al., 2017; Weible et al., 2009; Yadav et al., 2022). These diverse functions are supported by inputs from various sources including retrosplenial cortex (RSC), amygdala, thalamic nuclei, ventral tegmental area (VTA) and hippocampus (Cenquizca et al., 2007; Hoover & Vertes, 2007; Song et al., 2024). In turn, ACC projects to various cortical and subcortical regions, including the RSC, motor cortex, dorsal striatum, amygdala, thalamic nuclei, and VTA (Domesick, 1969; Gabbott et al., 2005; Gămănuţ et al., 2018; Jhang et al., 2018; Rajasethupathy et al., 2015; Song et al., 2024).

Given these, we first addressed whether ACC neurons constitute a homogeneous population, or if their functional properties vary based on their projection targets. We monitored the activity of distinct subclasses of ACC neurons projecting to specific brain regions while animals navigated through a virtual environment, which contained one salient location where reward was delivered. Spatial maps each encoding different environmental features co-exist among these neurons. Among them, RSC dysgranular zone (RSCd)-projecting neurons displayed a high concentration of place fields around the reward location while their activity was context specific. In a different context, they lost their reward specific response. Optogenetic activation of RSCd-projecting neurons at a specific location increased the number of hippocampal neurons activated at that specific location, whereas chemogenetic inhibition prevented the formation of the hippocampal place cell map *per se*. Additionally, optogenetic activation also decreased locomotion speed, mimicking the natural behavior of expectation to receive a reward. Our results demonstrate that ACC integrates both contextual and saliency information and specifically conveys the location of salient environmental features to the hippocampus in top-down fashion to update the spatial representation. This stands in contrast to the traditional bottom-up inputs from entorhinal cortex (EC) & CA3 to CA1 (Middleton & McHugh, 2016; Zutshi et al., 2022), which are not required for CA1 place cell formation. Our results challenge the canonical view of memory consolidation theory which posits a unidirectional flow of information from the hippocampus to the neocortex (Frankland et al, 2005) and instead highlights the critical role of bidirectional interactions between the hippocampus and neocortex in the formation, maintenance, and updating of hippocampal place cell maps.

## RESULTS

### Distinct classes of ACC neurons project to dmSTR, RSCg, and RSCd

To reveal a direct top-down projection from ACC to the hippocampus, we injected a retrograde AAV vector to the dorsal hippocampus. Despite previous reports of functional connectivity between the structures (Corcoran, et al., 2016; Yadav et al., 2022), our data did not reveal any strong direct projection (Fig. S1A, B), consistent with other studies (Bian et al., 2019; Oh et al., 2014; Rajasethupathy et al., 2015). We therefore decided to focus on potential indirect pathways which through an intermediary region could connect the two structures. RSC was identified as a prime candidate since it bi-directionally connects with both ACC and the hippocampus via parahippocampal regions, thereby functionally bridging these two structures (Corcoran et al., 2016; Lee et al., 2023; Tsai et al., 2022; Vann et al., 2009). Anatomically, RSC can be divided into two subregions, the granular and dysgranular (also called agranular) zones (RSCg and RSCd) based on the extent of development of the inner granular layer (layer 4) (Garey, 1999; Sugar, 2011), both of which receive ACC projections (Hoover & Vertes, 2007). Additionally, since the dorsal medial striatum (dmSTR) contributes to decision making, sensation, and emotional control (Cohen et al., 2021; Heilbronner et al., 2016), ACC neurons projecting to this region were also targeted.

To determine if dmSTR, RSCg, and RSCd receive inputs from common or distinct ACC neuronal populations, retrograde AAV vectors encoding either GFP or tdTomato were injected into two different projection sites, dmSTR and RSCd, dmSTR and RSCg, or RSCd and RSCg (Fig. 1A, S1C, D). As a result, dmSTR-projecting neurons were found predominantly in layers 2/3 (64.2 ± 6.3%, n = 12 brain slices from 3 mice), with fewer in layer 5 (35.8 ± 6.3%), whereas RSCg- and RSCd-projecting neurons were more abundant in layer 5 (81.0 ± 7.1% for RSCg-projecting neurons and 62.4 ± 15.8% for RSCd-projecting neurons, n = 12 brain slices from 3 mice for each group) than in layers 2/3 (19.0 ± 7.1% for RSCg-projecting neurons and 37.6 ± 15.8% for RSCd-projecting neurons) (Fig. 1A, B). The majority of neurons in ACC were labeled with a single color, indicating that different neuronal subpopulations in ACC target dmSTR, RSCg, and RSCd (Fig. 1C). Irrespective of their layer distribution or downstream targets, most of the labeled neurons showed immunostaining with the anti-CaMKIIα antibody, with a minority displaying anti-GAD67 immunoreactivity indicating that they were predominantly excitatory neurons (Fig. 1D-F, n = 6 brain slices from 2 mice for each group).

**Figure 1.**
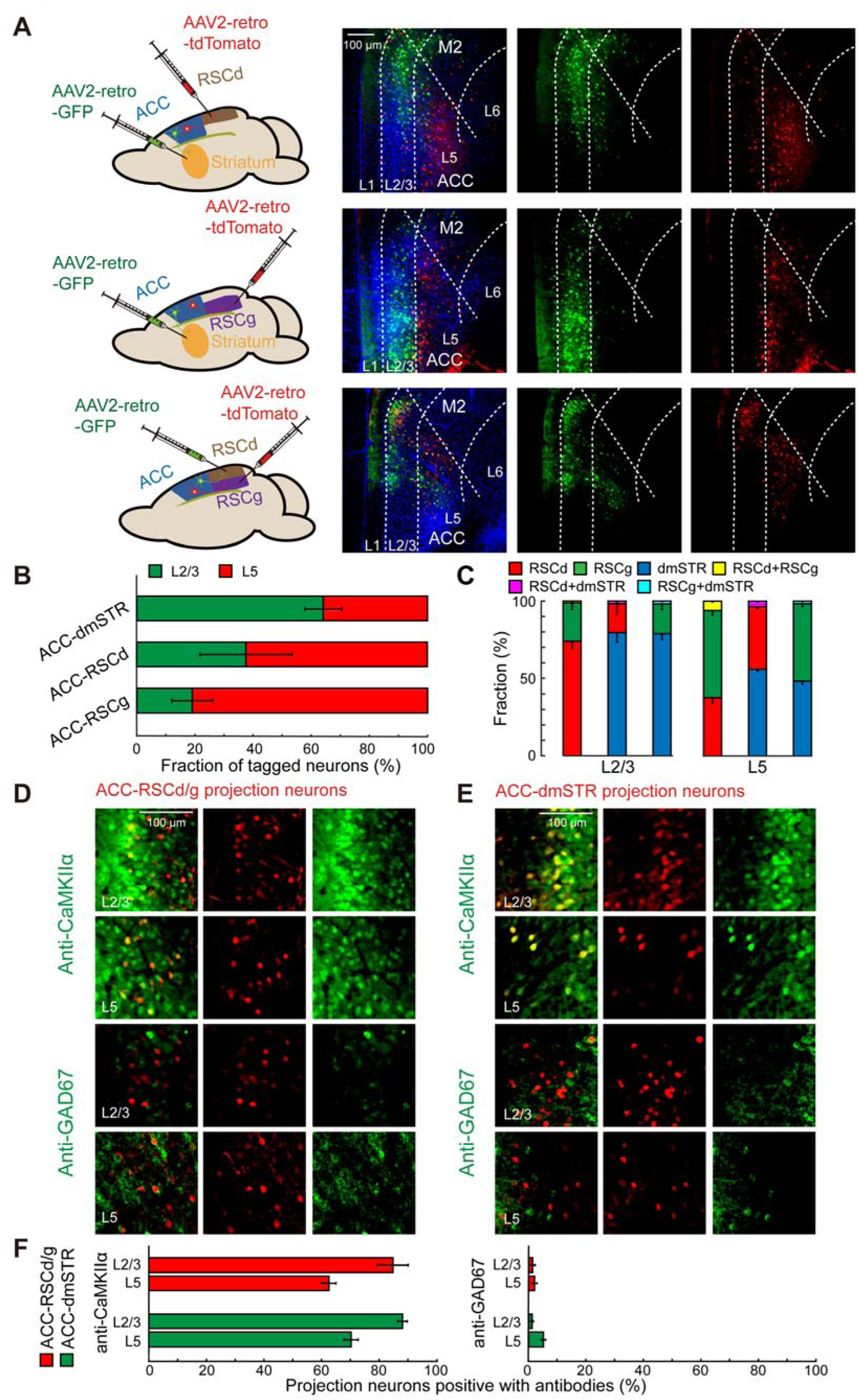
Distinct ACC neurons project to dmSTR, RSCg and RSCd. (A) Dual retrograde labeling of ACC neurons. Left: The injection sites of retrograde AAVs in RSCd, RSCg and dmSTR. Right: The distribution of fluorescently tagged neurons in the ACC. (B) The proportion of dmSTR-projecting, RSCd-projecting, RSCg-projecting ACC neurons located in ACC layer 2/3 or layer 5. (C) The fraction of ACC projection neurons which project to each area. (D) Example images of sections of ACC labeled with retrograde AAV injected in RSCd/g and immunostained with anti-CaMKIIα or anti-GAD67 antibodies. (E) Similar to (D) but injections of retrograde AAV were targeted to dmSTR. (F) The proportion of RSCd/g-projecting or dmSTR-projecting ACC neurons which were labeled with anti-CaMKIIα or anti-GAD67.

### Distinct activity patterns of ACC neuronal subpopulations during spatial navigation

We hypothesized that these distinct projections may convey different information to their downstream targets. To test this, a retrograde AAV vector encoding GCaMP6f was injected into either dmSTR, RSCg, or RSCd to monitor the activity of each type of projection neurons in ACC (Fig. 2A). A right-angle prism was inserted into the brain so that the vertical surface of the prism faced the median surface of the left ACC and the orthogonal surface faced upward (Low et al., 2014). After recovery, mice were head-fixed and habituated to running in virtual reality (VR) environments (Fig. 2B) without spatial cues, until mice ran more than 4000 cm in 20 min, which typically took 3-6 days. Following this, mice were exposed to two different VR tracks with distinct shapes (linear versus square) and wall patterns (Fig. 2C), while ACC layer 2/3 neuronal activity was monitored through the prism using a two-photon microscope. When mice reached the end or corner, the view of the track automatically turned, and the mouse resumed running in the new direction. Mice received a drop of water as a reward when they passed a fixed location in both environments. Each experimental day consisted of a linear track session followed by a square track session. As mice became more familiar with the task, they decreased their running velocity and began licking as they approached the reward location, demonstrating that they had associated a specific location with reward delivery (Fig. 2D, S2A-G).

**Figure 2.**
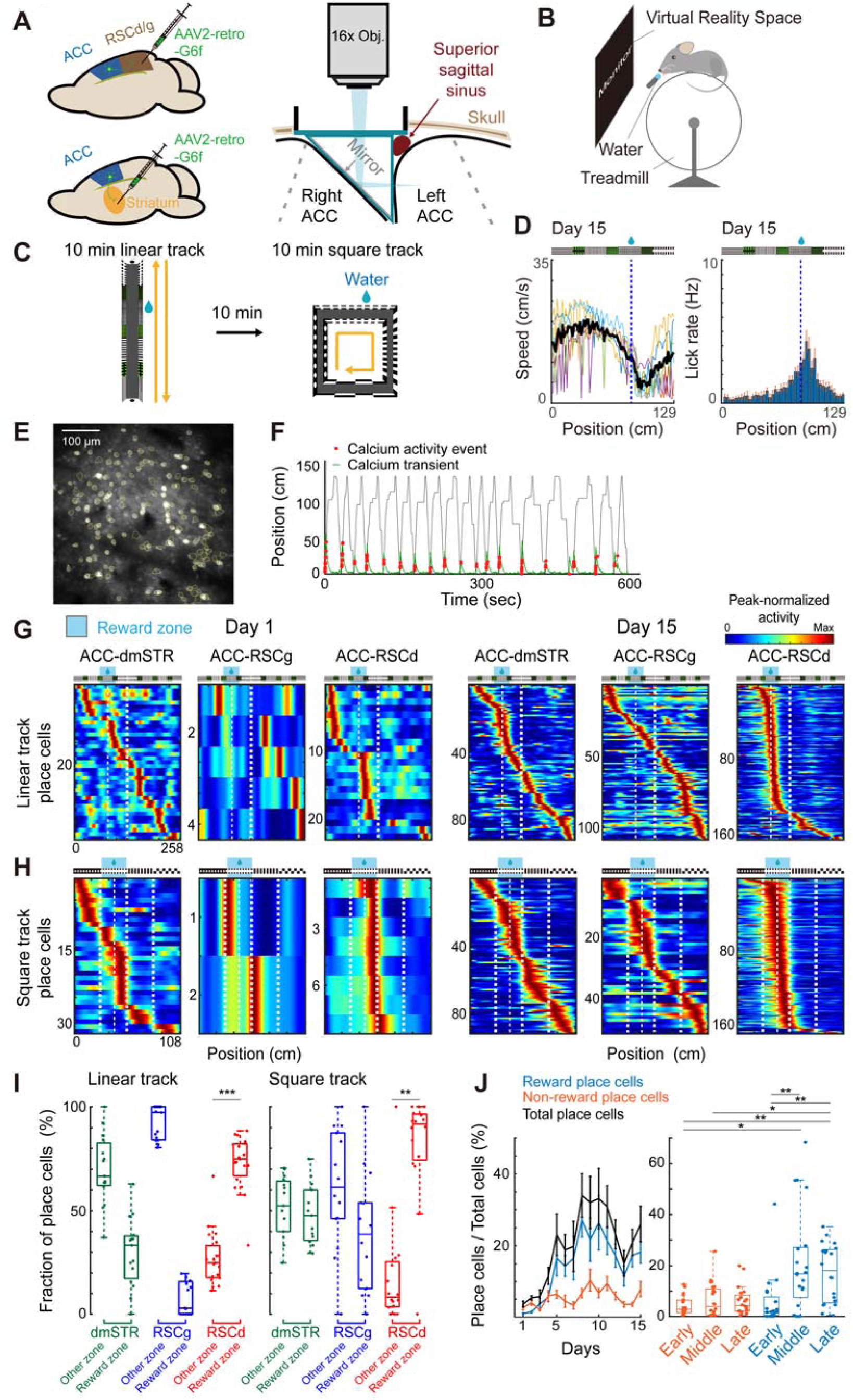
Different spatial maps are formed in dmSTR-, RSCg-, and RSCd-projecting ACC neurons. (A) Left: The injection sites of AAV2-retro-CaMKIIα-GCaMP6f. Right: Position of optical prism, depicting insertion into the right hemisphere and allowing for calcium imaging from the left ACC. (B) Virtual reality (VR) system for spatial navigation tasks in mice. Head fixed mice were placed on a treadmill and ran in a VR environment displayed on a monitor synchronized with the treadmill. A water spout was placed in front of the animal’s mouth, to allow delivery of water rewards. (C) The experimental protocol for VR tasks. (D) Mouse running speed and lick rate in the linear track on day 15. (E) Example of calcium imaging from ACC, with yellow contour lines depicting detected neurons. (F) Example of place cell activity in the VR task. The black line indicates mouse position, the green line indicates the calcium transient, with the red dots showing the location of calcium activity events. (G, H) The spatial activity maps of dmSTR-, RSCg- and RSCd-projecting place cells in the linear (G) and square tracks (H). Left: Day 1. Right: Day 15. (I) Left: Proportion of place cells with peak activity in the reward or non-reward zones for dmSTR-, RSCg- and RSCd-projecting neurons on the linear track. Right: The same for the square track reward zone (*** p < 0.001, ** p < 0.01). (J) Left: Proportion of reward and non-reward place cells among all RSCd-projecting neurons across days on the linear track. Right: Group data of reward and non-reward place cells proportions in early (days 1-5), middle (days 6-10) and late (days 11-15) sessions (*** p < 0.001, ** p < 0.01 and * p < 0.05).

On the first exposure to the environment, while dmSTR-projecting neurons showed a significant proportion of place cells (5.7 ± 1.3% of total 200 ± 29 neurons on the linear track and 4.7 ± 1.1% of total 201 ± 24 neurons on the square track, N = 3 mice), only a limited number of RSCg-projecting neurons displayed spatial specific activity (1.3 ± 0.4% of total 122 ± 29 neurons on the linear track and 0.6 ± 0.2% of total 116 ± 24 neurons on the square track, N = 3 mice) (Fig. 2E-H, S3). RSCd-projecting place cells comprised an intermediate proportion (3.6 ± 0.8% of total 198 ± 34 neurons on the linear track and 0.6 ± 0.2% of total 246 ± 50 neurons on the square track, N = 4 mice). Though the proportion of place cells increased over subsequent days and plateaued between day 5-10 in all three types of projection neurons (Fig. 2J, S3), this increase was not the result of changes in locomotion speed, which although is known to positively correlate with hippocampal place cell numbers (Sato et al., 2020), was comparable across days (Fig. S2H). Therefore, the increase was the result of repeated exposure to the same environment over days and increased familiarity (Fig. S2A-G).

There was a notable difference in the location preference of place cells among these projection neurons. Whereas dmSTR- and RSCg-projecting neurons represented the entire track, RSCd-projecting neurons activated almost exclusively around the reward location (Fig. 2G-J), indicating that depending on the downstream target, ACC conveys different information. Among the RSCd-projecting neurons, the number of place cells in the reward zone increased over days, while those in the non-reward zone remained comparable (Fig. 2J. Early, days 1-5; middle, days 6-10; late, days 11-15.). During late sessions, of all RSCd-projecting place cells, 73.0 ± 3.0% and 81.6 ± 6.5% activated near the reward location in the linear and square tracks respectively (displaying peak activity ±19.4 cm from the reward location in the linear track and ±13.5 cm in square track, chance level 15.0% and 25.0% respectively, assuming an even distribution along total lengths of 258 and 108 cm. Fig 2I. p < 0.001 for linear track and p *=* 0.003 for square track, Wilcoxon one-tailed signed rank test). In contrast, only 29.9 ± 4.3% and 48.1 ± 2.7% of dmSTR-projecting and 7.8 ± 2.1% and 39.0 ± 7.8% of RSCg-projecting cells showed spatial specific activity near the reward delivery site on the linear and square tracks respectively (Fig 2I). These results indicate that while RSCg- and dmSTR-projecting neurons convey information of the animals’ location within an environment, RSCd-projecting neurons specifically convey whether the animal is at the reward location or not.

### ACC reward place cells are specific to a context

To test whether the activity of RSCd-projecting neurons exhibit context specificity, we compared the activity of the same neurons in the linear and square tracks. The majority of cells activated at the reward location in one of the contexts, but not in another (Fig. 3A, B). While the proportion of place cells activity near the reward location increased over days, the proportion that activated in both tracks remained the minority (Fig. 2J, 3A-C). Among all RSCd-projecting place cells, 50.4 ± 7.7% and 37.0 ± 6.1% showed reward specific activity exclusively in the linear and square tracks respectively, while 12.6 ± 1.8% activated on both tracks in late sessions (N = 4 mice, Fig. 3A-C). This demonstrates that the RSCd-projecting neurons represent not just receipt of a reward, but also contextual information relating to the environment. We hereafter refer to RSCd-projecting neurons which preferentially activate around reward locations as reward place cells.

**Figure 3.**
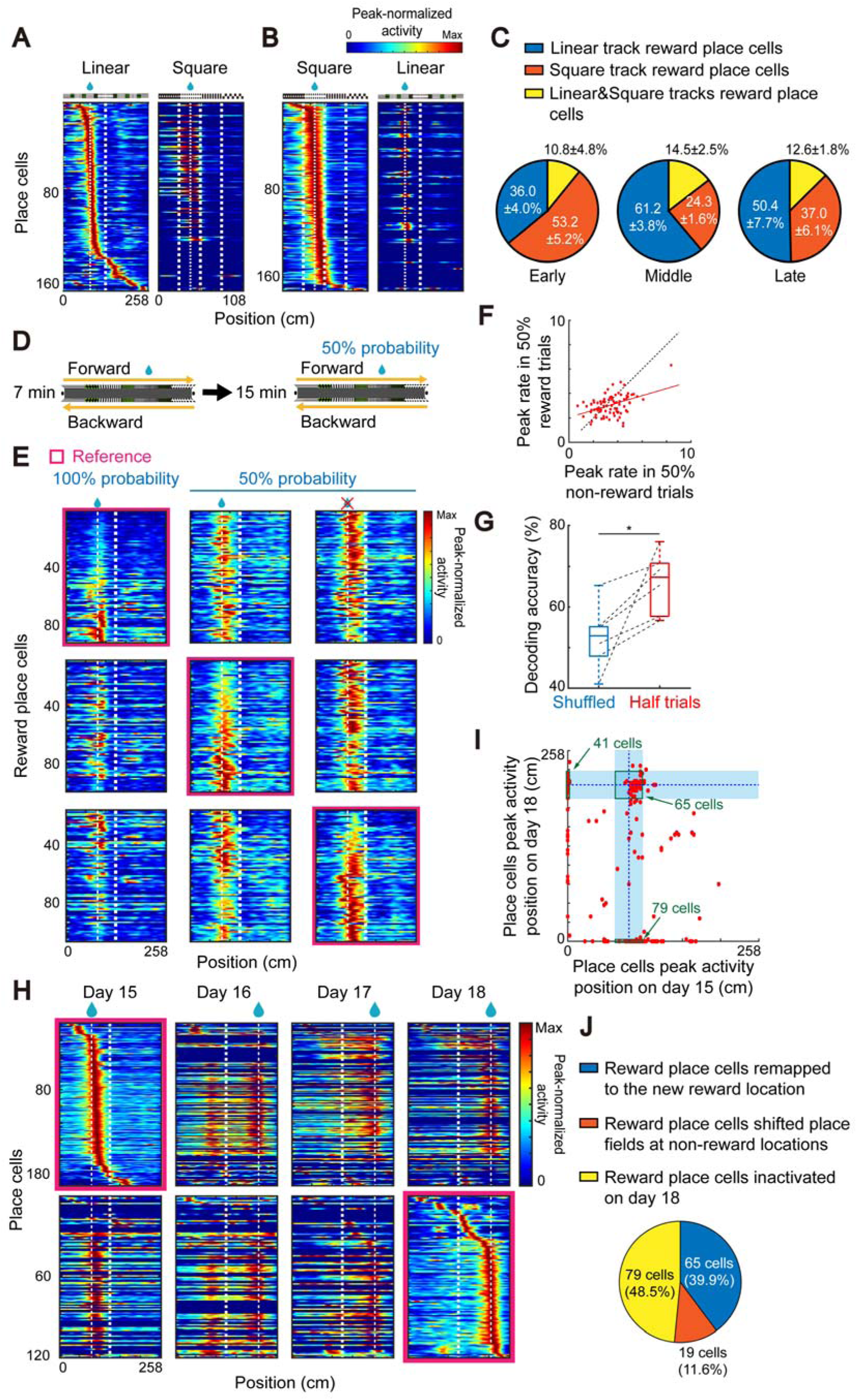
Context selectivity of RSCd-projecting reward place cells. (A, B) The calcium activity maps of place cells detected in the linear track and their activity in the square track (A), or vice versa (B). (C) The proportion of reward place cells in the linear, square or both tracks in early (days 1-5), middle (days 6-10) and late (days 11-15) sessions. (D) The experimental protocol for the reward omission task. (E) Calcium activity maps of reward place cells during the first 7 min trials (100% reward condition, left), followed by 15 min of the 50% random reward condition: rewarded trials (middle) and non-rewarded (right) trials. From top to bottom, the same set of neurons are sorted by activity in the 100% rewarded, 50% rewarded, and 50% non-rewarded trials, respectively. (F) The peak activity rate of each reward place cell in reward and no reward trials during the 50% random reward protocol. (G) Decoding of the presence or absence of reward using neuronal activity, by a logistic regression model of peak activity rate within the reward zone. Randomly shuffled activity of the same neuronal population was used to calculate significance (* p < 0.05). (H) Remapping of place cell activity maps in the reward shifting task. Reward location was shifted from day 16. Top: cells in each activity map sorted by place field position on day 15. Bottom: same as top but sorted by day 18. (I) The peak activity position of the same place cells on days 15 and 18, dots on the x and y axes indicate place cells which were detected only on days 15 and 18, respectively. (J) The proportion of reward place cells on day 15 that either remapped to the reward location, shifted to non-reward locations or became inactive on day 18.

To test whether the activity of reward place cells depends on reward receipt itself, or on prior reward experience two experimental designs were used. In the first experiment, animals first underwent a 15-day training protocol to familiarize them with the reward task. On day 16, they received rewards at 100% probability during the initial 7 minutes, after which for the following 15 minutes, rewards were delivered randomly at 50% probability (Fig. 3D). During the 50% probability trials, neurons continued to activate, with some showing enhanced activity in non-rewarded laps versus rewarded laps, potentially indicating that attention to unexpected events may drive increases in activity (Fig. 3E, F. p *=* 0.003, 50% reward trials compared with 50% non-rewarded trials by one sample *t*-test). We next attempted to test if it was possible to decode the presence or absence of reward based on the peak activity of these neurons within reward zone. Using the activity of RSCd-projecting neurons, a logistic linear classifier was able to decode the presence or absence of reward significantly better than shuffled data (Fig. 3G, p *=* 0.02, Wilcoxon one-tailed signed rank, n = 6 sessions, N = 3 mice). Overall, these data demonstrate that by modulating their activity, RSCd-projecting neurons in addition to coding reward location, also convey the presence or absence of reward in a specific context to downstream regions.

The second paradigm was used to probe how reward place cells respond to changes in reward location. As before, animals were trained for 15 days with the majority (89.6% of reward place cells, n = 144 neurons, N = 2 mice) of RSCd-projecting neurons activity tuned to the forward reward location (Fig. S4B). On day 16, animals received rewards randomly, either at the original forward location, or at a new backward location with 50% probability (Fig. S4A). Despite the change in reward delivery, the majority of reward place cells (96.9%) continued to activate in the original forward location even on reward omission trials, indicating that these neurons retain information encoding the location of the original reward (Fig. S4B, C). In addition, 52.4% of neurons activated in response to reward delivery in the new backward location. The activity at the new location was smaller and occurred only after reward delivery. Further, they displayed peak activity later in the forward direction than neurons that didn’t respond in the backward direction, indicating that these neurons activate in response to receipt itself, but not to expectation of the reward, unlike neurons in the forward direction whose activity begins as the animal approaches the reward (Fig. S4B). Together, these data indicate that reward place cells can retain previous reward locations even in the absence of reward itself but can still acutely remap to encode new reward locations.

### Remapping of reward place cells by chronic reward relocation

To understand how reward place cells respond to more chronic changes in the reward location, after 15 days of training with forward location, reward delivery was completely switched to a backward reward location for an additional 3 days (Fig. 3H). Of the reward place cells identified on day 15, when the reward was moved to the new location there was an overall decrease in the fraction of place cells on day 16, despite the total number of neurons remaining similar (18.6 ± 10.7% of total 220 ± 59 neurons on day 15 to 6.8 ± 3.2% of total 193 ± 63 neurons on day 16, N = 3 mice). Although the majority (63.2%) of reward place cells lost location specificity on day16, the remaining (36.8%) neurons began to activate at the new reward location, while retaining an additional field at the original reward location. By day 18, 39.9% of the original reward place cells from day 15 had remapped to the new reward location, 11.6% developed place fields at non-reward locations, with the remaining 48.5% becoming inactive (Fig. 3I, J). These data demonstrate that the activity of reward place cells is flexible and can track chronic changes in reward location.

### Activity of ACC-RSCd neurons is required for reward receiving behavior

Since environmental saliency has been shown to alter animal behavior (Engelhard et al., 2019; Sato et al., 2020), we hypothesized that the activity of these neurons might be important for inducing behavioral responses to saliency. To test this, AAV9-hSyn-DIO-hM4D(Gi)-mCherry and AAV2-retro-CaMKIIα-Cre were injected bilaterally into ACC and RSCd, respectively, allowing for selective chemogenetic silencing of RSCd-projecting neurons (Fig. 4A). Thirty minutes prior to the VR task, clozapine N-oxide (CNO) was administered intraperitoneally (*i.p.*) for 10 consecutive training days, after which training continued for a further 10 days in the absence of CNO (Fig. 4B). In the presence of CNO treatment, mice did not show the characteristic anticipatory behaviors associated with an upcoming reward observed in controls, such as reduction in running velocity and increased rate of licking (Fig. 2D, 4C. N = 4 mice for each group). When CNO was withdrawn however, the anticipatory behaviors became apparent (Fig. 4D), indicating that RSCd-projecting neurons are required for inducing saliency associated behaviors.

**Figure 4.**
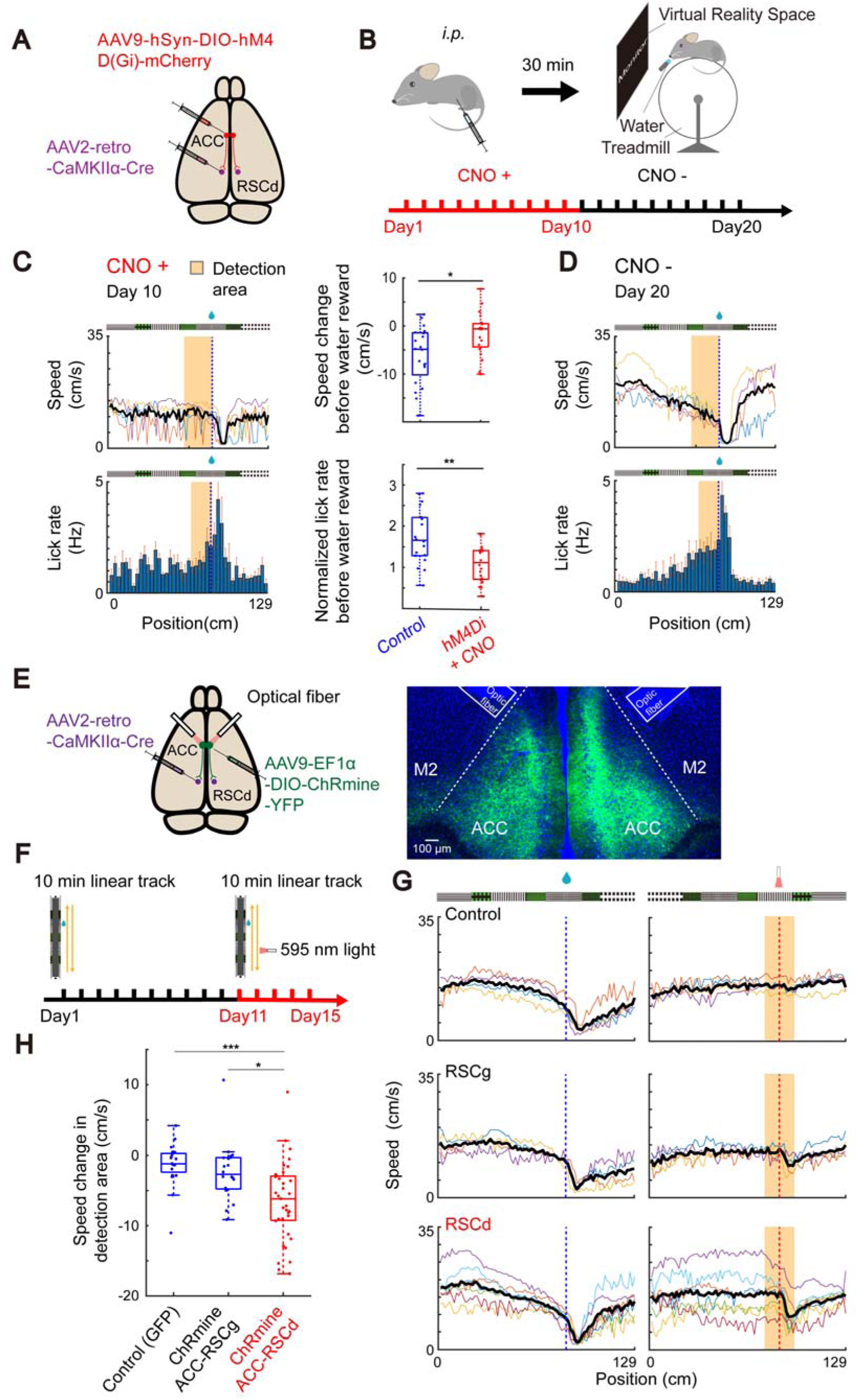
Activity of RSCd-projecting neurons is required for behavior associated with reward receipt. (A) The injection sites of AAV2-retro-CaMKIIα-Cre and AAV9-hSyn-DIO-hM4D(Gi)-mCherry. (B) The experimental protocol for RSCd-projecting neurons chemogenetic inhibition behavioral tasks. (C) Left: Running speed and lick rate in the reward direction on the last day of CNO treatment (day 10). Colored lines represent the data of individual mice, black line shows the mean. Right: The difference of lick rate, normalized by the mean lick rate in the whole linear track and running speed prior to reward location in the hM4Di+CNO (days 5-10) and control (days 5-10) groups (** p < 0.01 and * p < 0.05). (D) Same as (C) left but at day 20. (E) Left: Injection sites of AAV2-retro-CaMKIIα-Cre and AAV9-EF1α-DIO-ChRmine-YFP. Right: Confocal image showing ChRmine-YFP expression in ACC and the optical fiber tracks, scale bar represents 100 μm. (F) The experimental protocol for RSCd-projecting neuron optogenetic activation during behavioral tasks. (G) Average running speed of mice in the reward and light stimulation directions. Colored lines represent the average running speed from day 11-15 for individual mice. Bold black lines represent the average of all mice. ChRmine was expressed in either RSCg-projecting or RSCd-projecting ACC neurons. GFP was expressed in RSCd-projecting ACC neurons in control mice. (H) Change in running speed in the light stimulation zone (±12.4 cm from the light stimulation location, each dot represents the datum of a single mouse from one experimental day. *** p < 0.001, ** p < 0.01 and * p < 0.05).

To test if activation of RSCd-projecting neurons alone was sufficient to elicit the behavioral responses, the light-activated cation channel ChRmine was specifically expressed in RSCd-projecting neurons by injecting AAV9-EF1α-DIO-ChRmine-YFP and AAV2-retro-CaMKIIα-Cre into the ACC and RSCd respectively (Fig. 4E). In order to confirm that neurons could be activated sufficiently using this method, a cohort of mice underwent unilateral illumination in their home cages, followed by quantification of c-Fos expression in ACC. The number of c-Fos immuno-positive cells on the optically stimulated side was significantly higher than in the opposite unstimulated side, consistent with successful optical activation of RSCd-projecting neurons (Fig. S5A, B. p *=* 0.002, Wilcoxon one-tailed signed rank test. n = 12 brain slices from 3 mice).

Next, mice expressing ChRmine bilaterally were trained for 10 days in the VR linear track receiving reward in the forward direction. For the following 5 days, mice ran on the same track with an additional optical stimulation in the backward direction at a fixed location (Fig. 4F). Light delivery triggered an immediate reduction in animal running speed, that was specific to the mice (N = 7 mice) expressing ChRmine in RSCd-projecting neurons, but not in light-stimulated control mice (N = 4 mice) expressing GFP or mice expressing ChRmine in RSCg-projecting neurons (N = 4 mice) (Fig. 4G, H. p < 0.001 for RSCd versus control and p *=* 0.018 for RSCd versus RSCg, Kruskal-Wallis test with post-hoc test). These results demonstrate that RSCd-projecting neurons can modulate running speed and, coupled with the finding that light stimulation did not induce licking, is consistent with the notion that this RSCd-projecting pathway may be used as a route to convey saliency, rather than specifically reward location (Fig. S5C).

### Activity of ACC-RSCd neurons shapes hippocampal spatial representations

Given that the modulation of activity of RSCd-projecting neurons partly mimics the animals’ behavioral response to reward, we reasoned that this would also likely lead to changes in the hippocampal representation of space. To test this, ChRmine and GCaMP6f were expressed in RSCd-projecting neurons and dorsal hippocampus, respectively, together with a GRIN lens implanted above the hippocampus to monitor neuronal activity (Fig. 5A). Optogenetic activation of RSCd-projecting neurons increased the activity of 7.6 ± 1.2% of hippocampal neurons as assessed by increases in fluorescence immediately after each pulse of light (Fig. S6A, B. n = 26 ± 5 neurons, N = 4 mice.).

**Figure 5.**
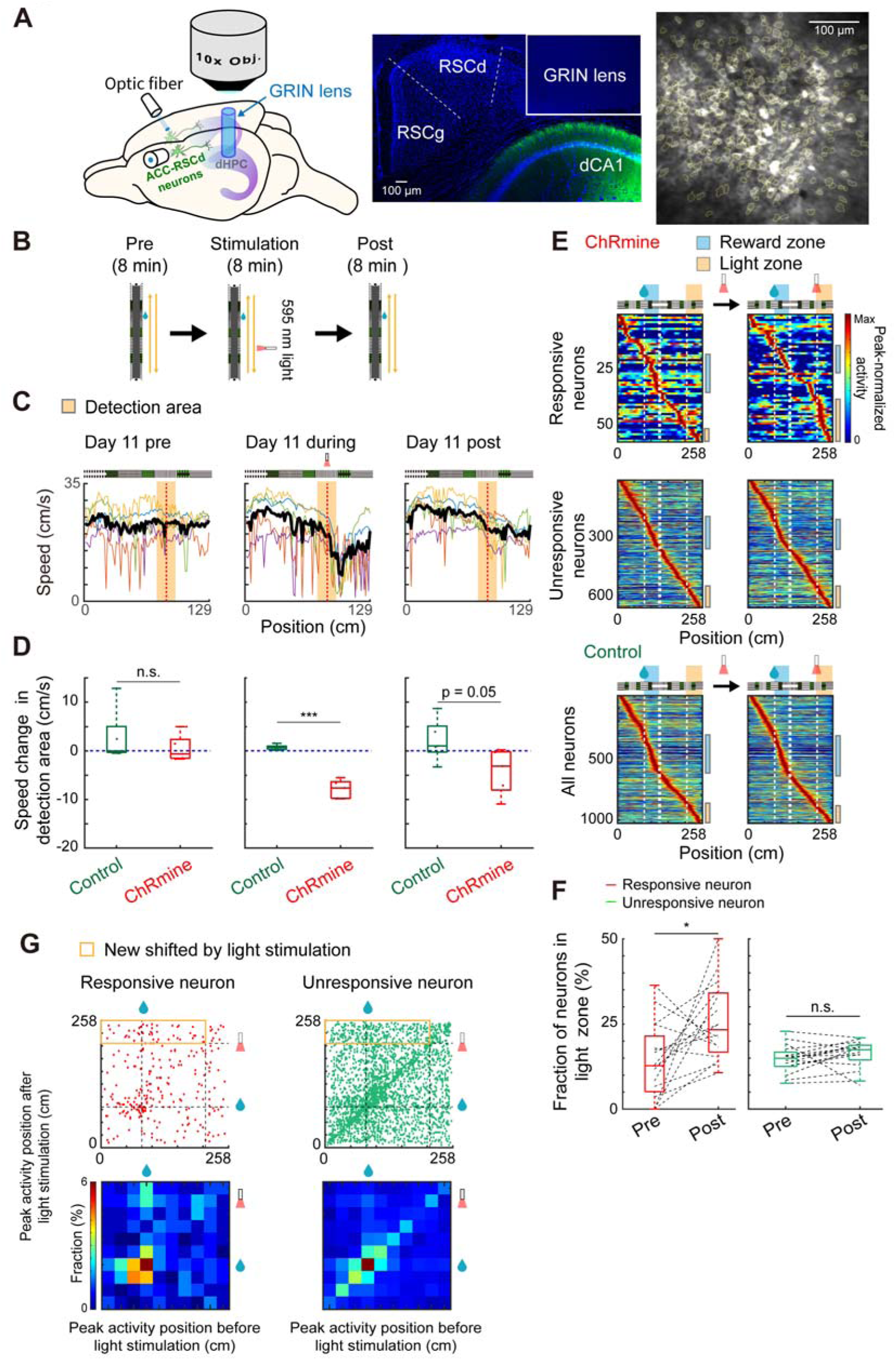
Activation of RSCd-projecting neurons acutely modifies hippocampal activity. (A) Left: The insertion location of optic fibers and GRIN lens for RSCd-projecting neuron activation and hippocampal imaging. Middle: A confocal image of a coronal section of a mouse brain showing the GRIN lens location. Right: Example of calcium imaging for dorsal hippocampus, with yellow contours showing detected neurons, scale bar represents 100 μm. (B) The experimental protocol of RSCd-projecting neuron activation and imaging. (C) The running speed of ChRmine expressing mice before, during, and after the optical stimulation of RSCd-projecting neurons. Colored lines represent the average running speed for individual mice. Bold black lines represent the average of all mice. (D) The speed change of control and ChRmine expressing mice in detection area before, during, and after the light stimulation (*** p < 0.001). (E) The activity maps of light-responsive and unresponsive neurons in ChRmine mice and all neurons in control mice before and after light stimulation. Blue and orange bars on right indicate neurons have peak activity in the reward and light-stimulation zones, respectively. (F) Fraction of neurons which have peak activity in the light stimulation zone before and after light stimulation in responsive and unresponsive neurons. (G) Top: The peak activity position of light-responsive and unresponsive neurons before and after light stimulation. Bottom: The proportion of responsive and unresponsive neurons which have peak activity in each location block. The area of top map was divided into 10 x 10 blocks.

Mice were trained as described previously for 10 days, after which on days 11 to 14, 24 min sessions were conducted consisting of; an 8 min pre-stimulation baseline period, followed by an 8 min stimulation period, where RSCd-projecting neurons were stimulated when mice ran through a zone without a reward and finally an 8 min post-stimulation period concluded each day (Fig. 5B). As before, light stimulation of RSCd-projecting neurons triggered a decrease in animal running velocity (Fig. 4G, 5C). This effect persisted through the post-stimulation session, albeit with a smaller reduction in speed (Fig. 5C, D. p = 0.03, Wilcoxon rank sum test. N = 5 mice for each group.), suggesting that RSCd-projecting neuron activation can induce neural changes that lead to persistent changes in behavior.

To identify an underlying mechanism, hippocampal neuronal activity was monitored before and after the light stimulation (Fig. 5E). Consistent with the lasting behavioral changes observed, the number of neurons active in the light stimulation zone increased in the post-stimulation period (Fig. 5E, F). This increase was observed only among light-responsive neurons, but not among unresponsive neurons, or in control animals not expressing ChRmine (p = 0.01, Wilcoxon signed rank test for responsive neurons, n = 16 sessions, N = 4 mice).

Tracking the peak activity position of hippocampal neurons across the pre- and post-stimulation periods, revealed a significant proportion of the light unresponsive neurons maintained a stable location selectivity (distributed along the y = x diagonal) despite the light stimulation (Fig. 5G). In contrast, a fraction of the light responsive neurons that activated outside the reward zone prior to light stimulation, were recruited to the light stimulation zone in the post-stimulation session (n = 269 responsive and n = 3136 unresponsive neurons, Fig. 5G), indicating that stimulation of RSCd-projecting neurons in ACC can acutely remap hippocampal place cells to signal salient locations in an environment.

### Activity of ACC-RSCd neurons can alter stability of the hippocampal map

To assess how long RSCd-projecting neurons artificial activation triggered hippocampal remapping persists, the distribution of place cells in pre-stimulation sessions across successive days were quantified. On day 11, prior to light stimulation, there was a significant enrichment of place cells near the reward location in the both ChRmine and control groups (1.6 ± 0.3-fold versus 1.4 ± 0.3-fold compared with an even distribution, N = 5 mice for each group), consistent with previous studies (Sato et al., 2020), which was not the case for the light-stimulation zone (0.8 ± 0.2-fold versus 0.8 ± 0.3-fold in ChRmine and control groups, Fig. 6A-C). After repeated light stimulation over days, there was a marked increase in the number of neurons representing the light-stimulation zone in ChRmine expressing versus control mice (1.1 ± 0.1-fold compared with an even distribution on day 14, control mice 0.6 ± 0.2-fold, p = 0.01, two-way ANOVA for days 12-14). Interestingly, this was associated with a decrease in the fraction of neurons representing the reward zone in the ChRmine versus control group (1.3 ± 0.3-fold versus 1.9 ± 0.6-fold on day 14 in ChRmine and control mice respectively, p = 0.03 for days 12-14, Fig. 6A-C).

**Figure 6.**
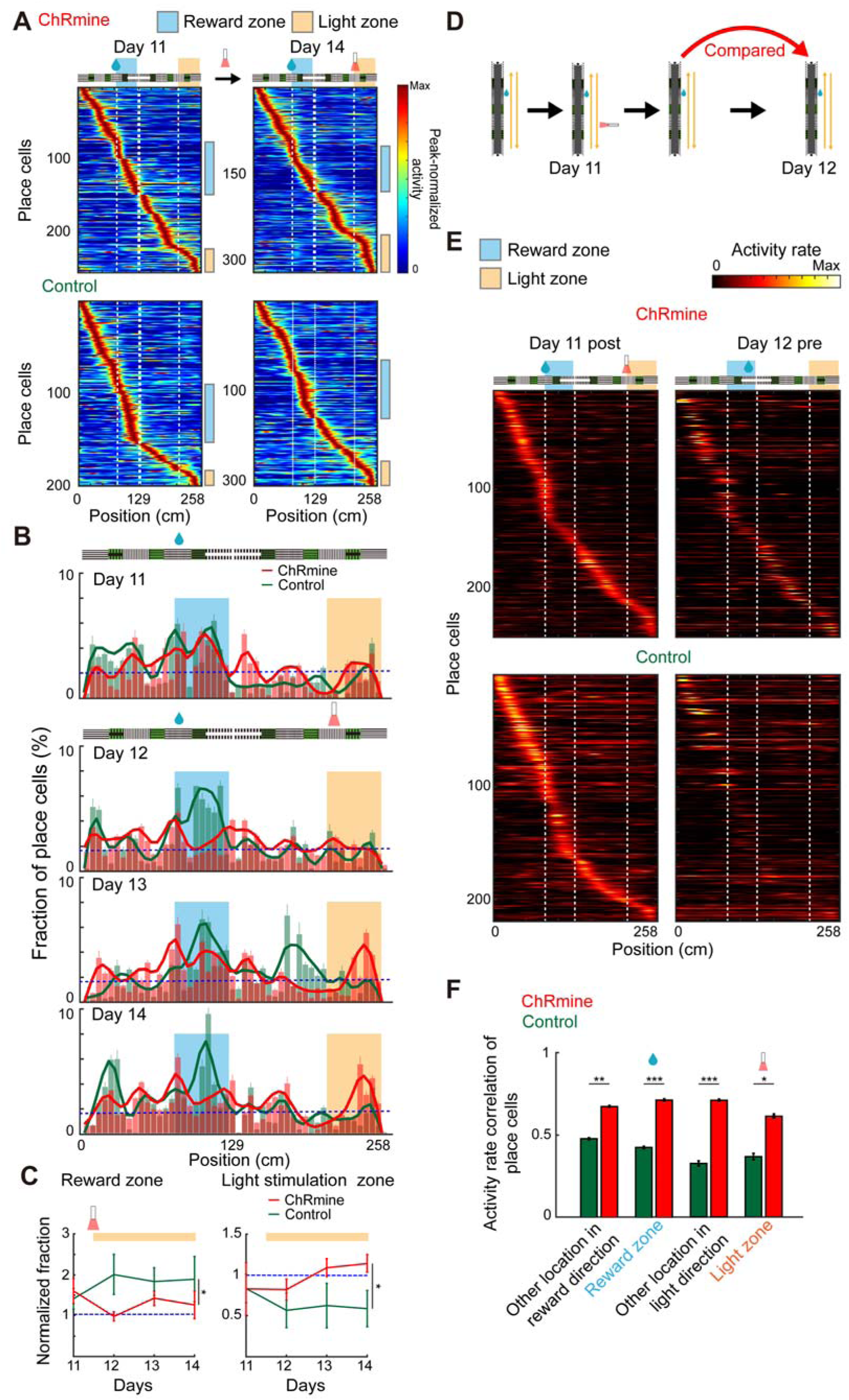
Activation of RSCd-projecting neurons locally enriches the hippocampus spatial representation. (A) The hippocampal spatial activity maps of mice expressing ChRmine in RSCd-projecting ACC neurons and stimulated as in Fig. 5B for four consecutive days on days 11 and 14. Hippocampal neurons were imaged on day 11, followed by activation of RSCd-projecting neurons. On day 12 and subsequent days, the same procedure was repeated, thus the data shown as Day 14 were obtained 24 hours after the last activation. Top: ChRmine group. Bottom: Control group. Blue and orange bars on right indicate place cells detected in the reward and light-stimulation zones, respectively. (B) Histogram (bars) and kernel density estimation (curves) showing the distribution of place cells detected in each spatial bin along the linear track. (C) Left: The fraction of dorsal hippocampal place cells in the reward zone (44 cm after reward location, normalized to an even distribution of place cells across the linear track). Right: The fraction in the light stimulation zone (44 cm after light stimulation location. * p < 0.05). (D) Protocol of optogenetic experiments to test the effects of RSCd-projecting neuron activation on place cell stability. (E) Activity maps of the day 11 post-stimulation session (left) and the activity of the same neurons in the day 12 pre-stimulation session (right) in control and ChRmine expressing neurons. (F) Activity correlation of place cells between day 11 post-and day 12 pre-stimulation sessions.

Our previous work demonstrated that environmental saliencies such as rewards or landmarks enhance the cellular representation of these locations, by stabilizing hippocampal place fields at the location (Sato et al., 2020). To establish if stimulation of RSCd-projecting neurons alters the stability of the hippocampal representation of space, the spatial maps of post-stimulation sessions were compared with those of pre-stimulation sessions from the following day (Fig. 6D). This revealed the maps across successive days were more correlated in stimulated versus control animals, indicating that activation of RSCd-projecting neurons stabilizes hippocampal spatial representations (Fig. 6E, F). Notably, this stabilization was not confined to the light stimulation zone but encompassed the entire track in both running directions (Fig. 6F, n = 250 & n = 219 place cells for the ChRmine and control groups respectively, p < 0.001 & p = 0.017 for the reward and light stimulation zones respectively, Wilcoxon rank sum test). Given that only a small fraction of hippocampal neurons is directly stimulated via the light-activation of RSCd-projecting ACC neurons, this indicates that the effects of light stimulation on place cell stability are likely spread to non-place cells as well. To confirm this, the stability of non-place cells was calculated separately, which revealed a higher degree of stability in the ChRmine group (Fig. S6C, n = 457 & n = 615 neurons for the ChRmine and control groups respectively, p < 0.001, Wilcoxon rank sum test). This indicates that while RSCd-projecting neurons directly recruit hippocampal neuronal activity in the light stimulation zone, they also act to increase the overall stability of the entire hippocampal population.

### Inhibition of ACC-RSCd neurons prevents the formation of the hippocampal place cell map but does not affect it once formed

Given the ability of the RSCd-projecting neurons to trigger changes in stability and broadcast salience to the hippocampus, to better understand their downstream effects we used a chemogenetic silencing strategy while simultaneously imaging hippocampal activity (Fig. 7A-C). Though inactivation of RSCd-projecting ACC neurons did not affect the number of neurons detected (Fig. 7D), it reduced the number of place cells in the hippocampus by impairing their spatial specific activity (Fig. 7B-E. N = 4 mice & N = 3 mice for hM4Di and control groups respectively). Following cessation of chemogenetic silencing, the number of place cells increased with additional training (Fig.7B, D). By contrast, if a pre-existing dorsal hippocampal spatial map was already formed following 10 days of training, inactivation of RSCd-projecting ACC neurons on day 11 had no effect (Fig. 7F, G). Together, these results demonstrate that the top-down signal from RSCd-projecting neurons is necessary for the initial formation of spatial maps in the hippocampus, but once established are no longer required for their maintenance.

**Figure 7.**
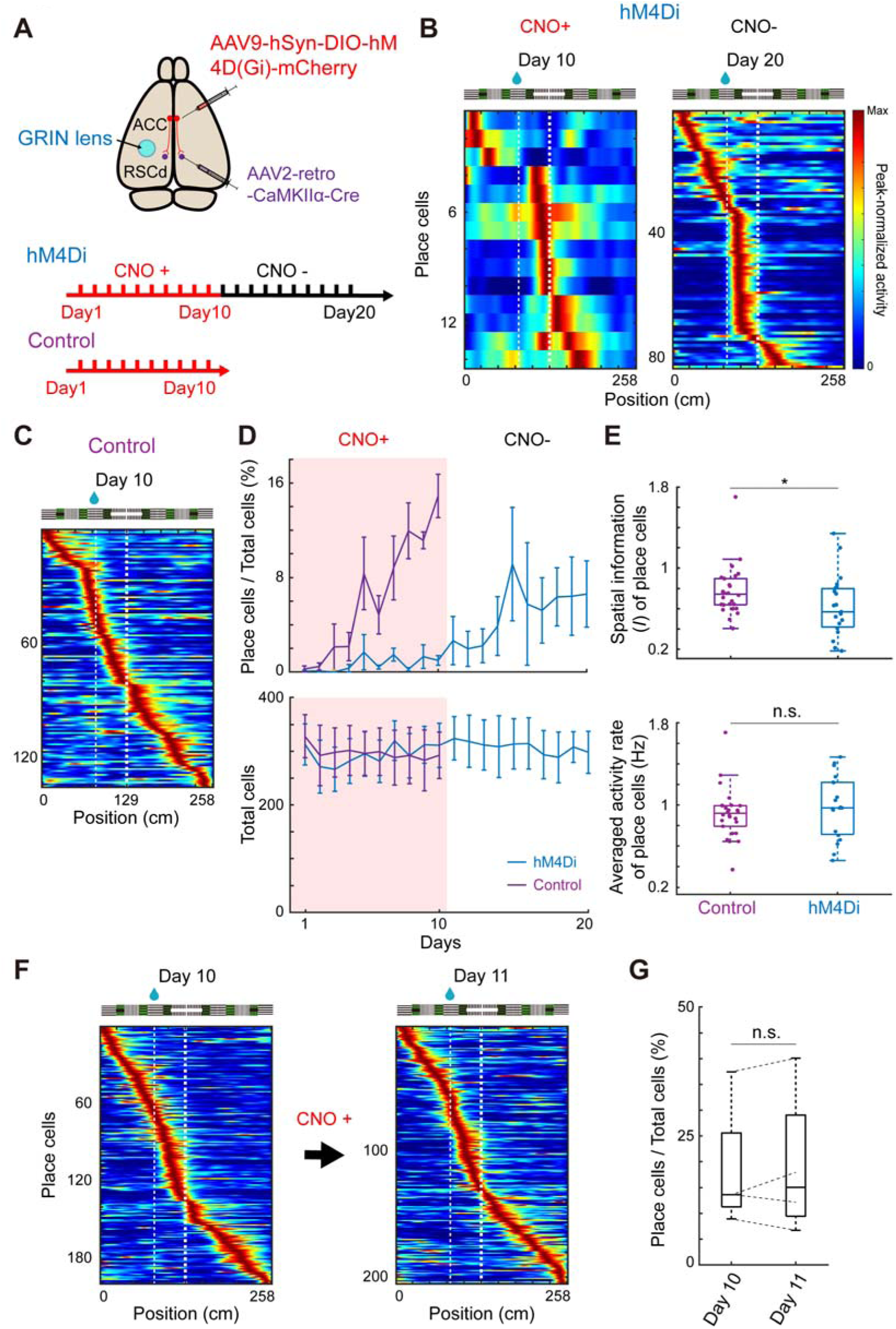
Inhibition of RSCd-projecting neurons disrupts dorsal hippocampus spatial map formation. (A) Top: the injection sites of AAV2-retro-CaMKIIα-Cre and AAV9-hSyn-DIO-hM4D(Gi)-mCherry. A GRIN lens was implanted directly above dorsal CA1. Bottom: Experimental schedule. (B) The activity maps of hippocampal place cells on the last day of ACC-RSCd inhibition by CNO (day 10) and 10 days later in the absence of CNO (day 20). (C) The activity map of hippocampal place cells on the last day of CNO treatment (day10) in control mice. (D) Top: The fractions of hippocampal place cells in the hM4Di expressing and control groups. Inhibition was only performed during the first 10 days. Bottom: The number of detected neurons in the hM4Di and control groups. (E) Top: The spatial information (*I*) of place cells in the hM4Di and control groups. Each dot represents one session in a mouse, no place cell detected sessions were excluded. Bottom: The averaged activity rate of place cells in the hM4Di and control groups (* p < 0.05). (F) The activity maps of dorsal hippocampal place cells in mice receiving CNO on day 11 after dorsal hippocampal spatial maps were already formed during 10-days of prior training. (G) The proportion of dorsal hippocampal place cells in mice shown in (F).

## DISCUSSION

Here we identified a specific top-down input from the ACC to the RSCd that becomes active exclusively at reward locations. Activity of these ACC neurons is involved in spatial navigation learning and is both required and sufficient for shaping the hippocampal saliency map.

Reflecting its diverse functions, ACC projects to various brain regions including the striatum, RSC, hippocampus, and thalamus (Domesick et.al, 1969; Gabbott et al., 2005). While prior studies have focused on these functions of the ACC, how information on environmental saliency is encoded, segregated, and routed to downstream brain regions, particularly at the single neuron level has been neglected. In this study we used retrograde tracing to separately label the ACC neurons that project to the dmSTR, RSCd, and RSCg and found distinct populations project to each region, with little overlap (Fig. 1), supporting the notion that information is segregated across ACC and routed to specific downstream targets.

Our results demonstrated that there were clear differences among the three classes of projection neuron. dmSTR-projecting and RSCg-projecting ACC neurons showed activity similar to hippocampal place cells, activated sequentially as animals traversed their environment and forming a spatial map which spanned the entire linear track. By contrast, the activity of RSCd-projecting ACC neurons was highly concentrated around reward locations (Fig. 2). Moreover, the multi-environment experiments demonstrate that these neurons become tuned to a specific context (Fig. 3). These results suggest that RSCd-projecting neurons encode rewards tied to a specific location, rather than producing a general response to any reward and thus, signal whether an animal is at a specific reward location or not. These results are the first to identify a novel class of principal neuron in a corticocortical pathway, that specifically processes information about reward location.

Optical activation of RSCd-projecting ACC neurons induced a reduction in animal velocity consistent with the behavioral response associated with expectation of an upcoming reward, that was absent when RSCg-projecting ACC neurons were stimulated (Fig. 4E-H). The effect persisted after the optical stimulation session indicating that the stimulation was sufficient to form a memory (Fig. 5C, D). The chemogenetic inhibition experiments demonstrate that the ACC-RSCd circuit is required for associating a specific location with a reward. In the absence of activity of these neurons, mice failed to form the characteristic behavioral signatures of reward-receipt, such as decreased velocity on approach to the reward and increased lick rates. Only after cessation of CNO administration, did mice begin to display behavior consistent with task learning (Fig. 4D). Though previous studies have implicated ACC in reward learning (Akam 2021; Teixeira et al., 2006), our results indicate that this function can be attributed to a subset of ACC neurons that project to RSCd.

Experimental evidence of how the spatial maps co-existing in both ACC and hippocampus influence each other, and how they are updated during behavior is sparse (Bota et al., 2021; Rajasethupathy et al., 2015; Kitamura et al., 2017). We demonstrated that activation of RSCd-projecting ACC neurons triggered activity in a subpopulation of hippocampal neurons. If stimulation of these neurons was coupled to a specific location on the track over the course of several days, there was a corresponding enrichment of hippocampal place cells at that location (Fig. 6). Crucially, while inhibition of the RSCd-projecting ACC neurons prevented the normal formation of a hippocampal place cell map, this was only the case if the manipulation was conducted during learning but had no noticeable effect once the animal already learnt the reward location (Fig. 7). Taken together, these findings demonstrate that the ACC exerts top-down control over the hippocampal spatial representation, via RSCd. Once reward information is routed to and stabilized in hippocampus, it persists and is no longer altered in the absence of ACC-RSCd input, thus leading to the retention of locations containing salient environmental cues, but when required this pathway can be activated to update the hippocampal map to signal salience at a new location.

Previous studies showed that artificially activated locus coeruleus input to dorsal hippocampus coupled with reward can also induce enrichment of hippocampal place cells near specific location (Kaufman et al., 2020). In addition, dopamine signals are known to play an important role in reward learning, and VTA neurons are their main source (Fields et al., 2007; Hyman 2006). VTA sends dopaminergic projections to multiple brain regions including the hippocampus, but the axon terminals are mainly located in the ventral hippocampus and extremely sparse in dorsal CA1 (Kempadoo et al., 2016; Takeuchi et al.,2016). In contrast to dorsal CA1, ACC receives strong projections from the VTA (Chen et al., 2024; Song et al.,2024) which is important for reward learning (Elston et al., 2018; Narita et al., 2010). Therefore, such input from VTA to ACC are also required for establishing reward-selective responses of RSCd-projecting ACC neurons. However, it should be noted that there is a clear difference between RSCd-projecting ACC neurons and VTA dopaminergic neurons. Unlike the dopaminergic neurons, which respond to a reward itself, or even to cues that lead to expectation of a reward (Cohen et al.,2012; Hu et al., 2016; Schultz et al., 1997), RSCd-projecting neuron activity is tied to a specific reward location and is significantly reduced if the reward location is moved. Our results suggest that ACC functions as a node where spatial recognition systems and reward systems converge.

The characterization and understanding of this top-down pathway, which specifically processes reward location information and how it exerts control over the hippocampal representation of space, broaden our understanding of how reward information is processed, stored and updated as required for spatial memories. Contrary to existing literature which points to hippocampus as the first structure to encode important changes as external conditions alter and send the information in a bottom-up fashion to association cortex (reviewed in Frankland et al., 2005), our data implicate ACC as the primary driver of this process. Specifically, we found that a top-down signal from a subclass of neuron in ACC is required for the initial formation of the hippocampal map, as well as updating it to incorporate novel salient locations. Intriguingly, once the hippocampal map is consolidated, it no longer requires input from ACC to persist. Overall, these results are unexpected from the canonical memory consolidation theory where information always flows in a bottom-up fashion. Also, this is in stark contrast with previous studies of blockade of the two main sources of input to CA1 (EC and CA3), which do not preclude the formation of place cells (Middleton & McHugh, 2016; Zutshi et al., 2022). Future studies will likely address both the contributions of deep layer neurons and target multiple ACC populations simultaneously to identify how the ACC place and reward maps are reconfigured as rewards are altered.

## ACKNOWLEDGMENTS

We thank Drs. Kenji Mizuseki and Thomas J. McHugh for valuable comments on the manuscript. This work was supported by Grant-in-Aid for Scientific Research JP21650080, JP16H01292, JP16H01438, JP16H02455, JP17K19631, JP18H05434, JP19H01010, and JP22H04981 from the MEXT, Japan, The Uehara Memorial Foundation, The Naito Foundation, Research Foundation for Opto-Science and Technology, Novartis Foundation, and The Takeda Science Foundation, HFSP Research Grant and RGP0020/2019, and JST CREST JPMJCR20E4 to Y.H., JST SPRING, Grant-in-Aid for Scientific Research JPMJSP2110 and JP22K20684 to Y.L., and Grant-in-Aid for Scientific Research Number JP21H00310 and JP23K24199 to K.M.

---

**Supplementary Figure 1.**
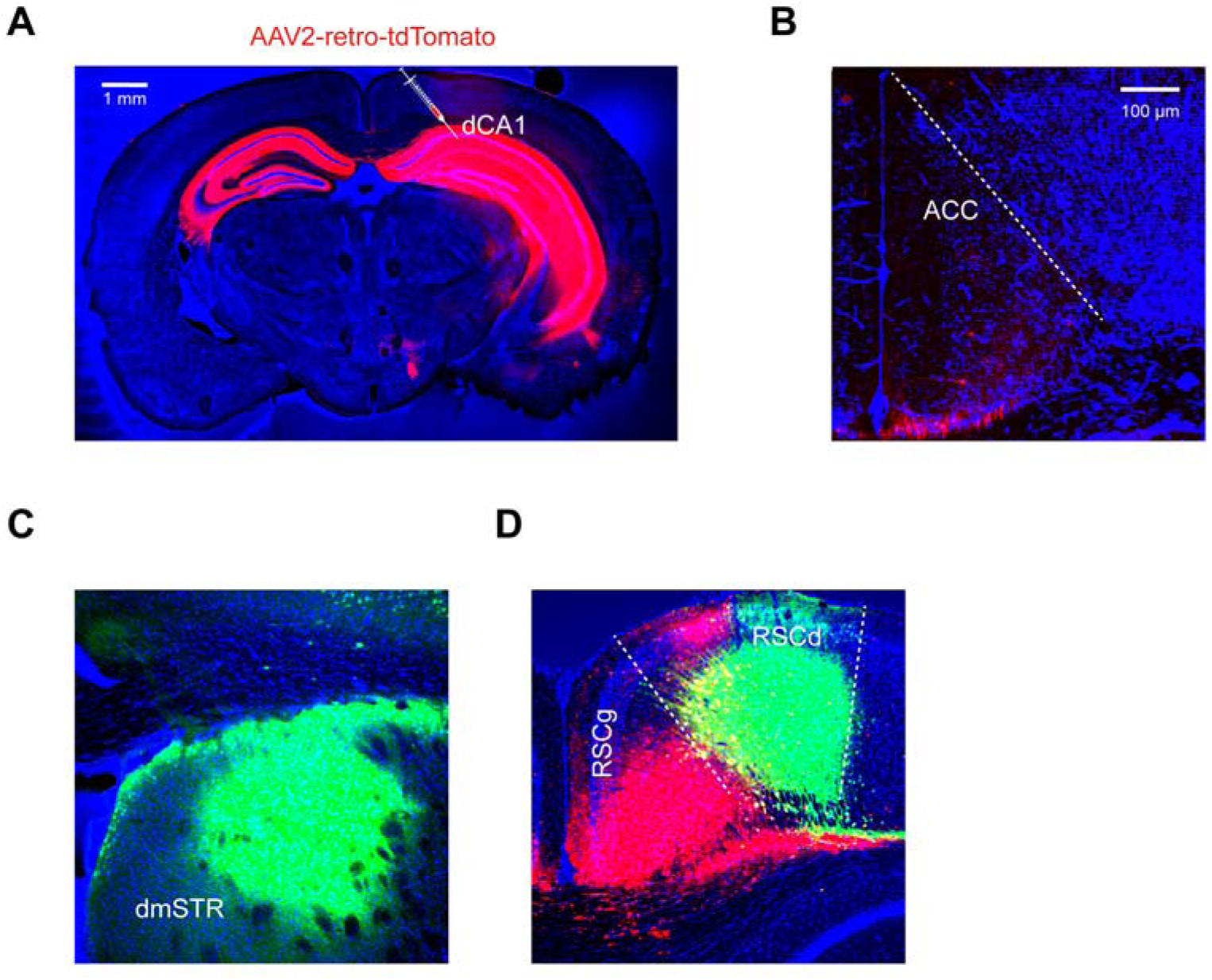
Retrograde labeling of ACC by retrograde AAVs. (A) An example of the injection site of AAV2-retro-tdTomato in dorsal hippocampus. (B) ACC section of the same animal showing that neurons projecting to dorsal hippocampus are scarce. (C) An example of the injection site of retrograde AAV into dmSTR. (D) An example of the injection sites of retrograde AAVs into RSCd or RSCg.

**Supplementary Figure 2.**
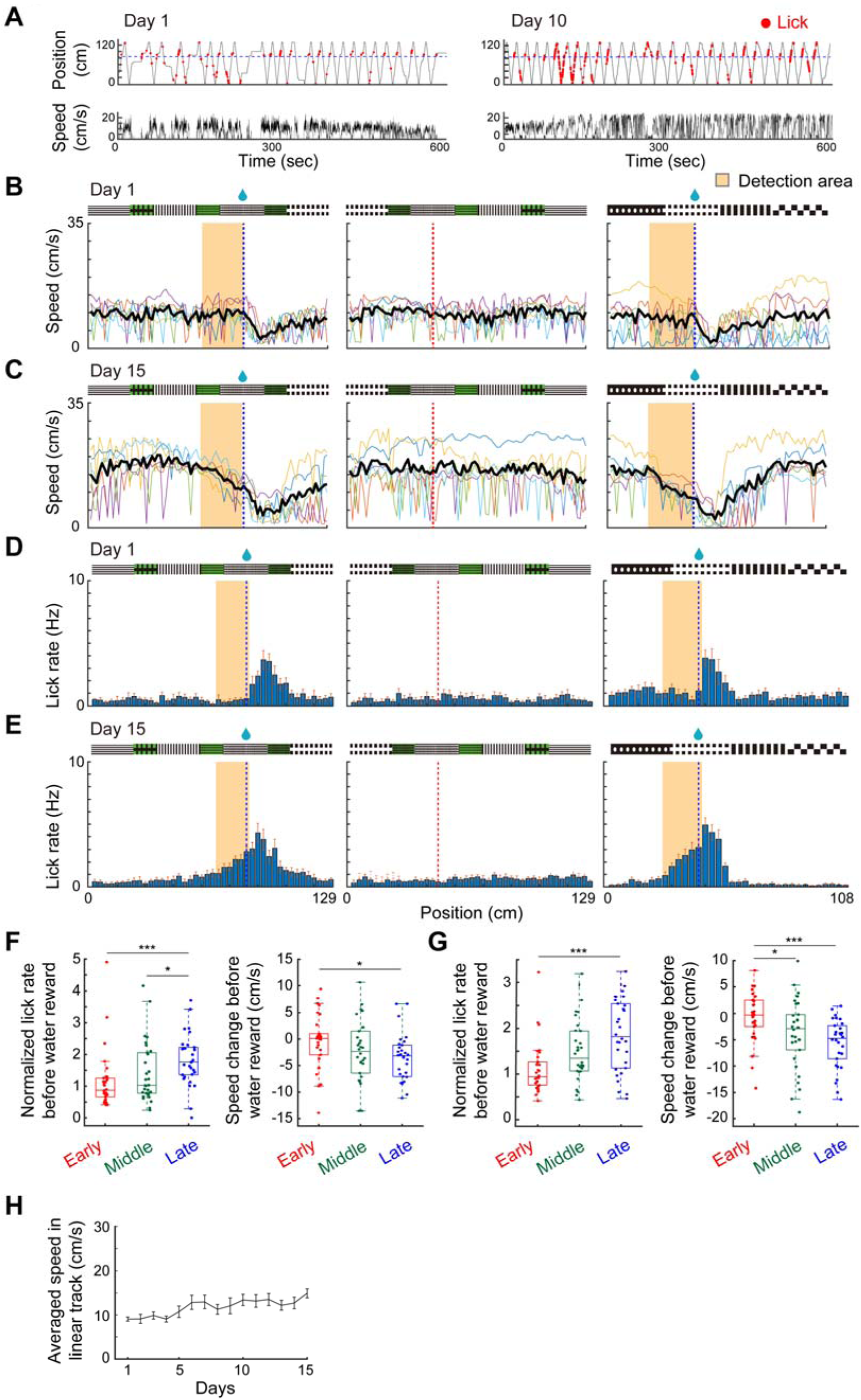
Mice behavior in spatial navigation learning tasks. (A) Examples of mouse behavior in day 1 and day 10 sessions. (B) Left: Running speed in the linear track reward direction on day 1. Middle: In the non-reward direction on day 1. Right: In the square track on day 1. Colored lines represent the average running speed for individual mice. Bold black lines represent the average of all mice. (C) Same as (B) but on day 15. (D) Left: Lick rate in the reward direction of the linear track on Day 1. Middle: Lick rate in the non-rewarded direction of the linear track on Day 1. Right: Lick rate in the square track on Day 1. (E) Same as (D) but on day 15. (F) Left: Lick rate normalized by the mean rate across the entire linear track. Right: The difference in running speed before reward location (0-10.5 cm prior to reward location) in the linear track (Early, day 1-5; Middle, day 6-10; Late, day 11-15). (G) Similar plot to (F) in the square track. (H) Mean running speed of mice in the linear track across days.

**Supplementary Figure 3.**
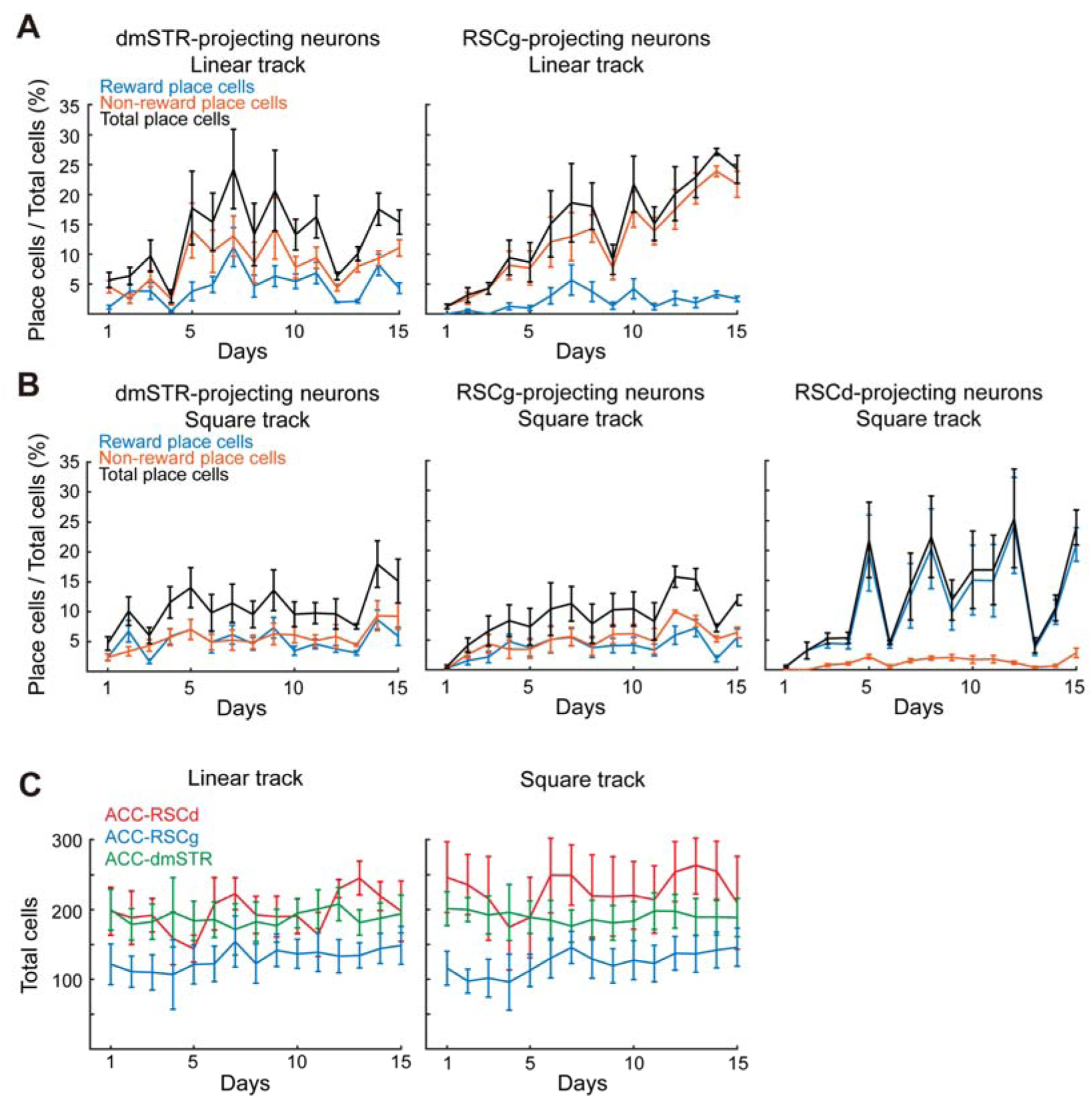
Proportion of detected cells across days. (A) Left: The proportion of reward and non-reward place cells among all dmSTR-projecting neurons across days in the linear track. Right: Same as left, but for RSCg-projecting neurons. (B) Left: The proportion of reward and non-reward place cells among all dmSTR-projecting neurons across days in the square track. Middle: Same as left, but for RSCg-projecting neurons. Right: Same as middle, but for RSCd-projecting neurons. (C) The total detected cells of all three types of projection neurons across days in the linear (left) and square tracks (right).

**Supplementary Figure 4.**
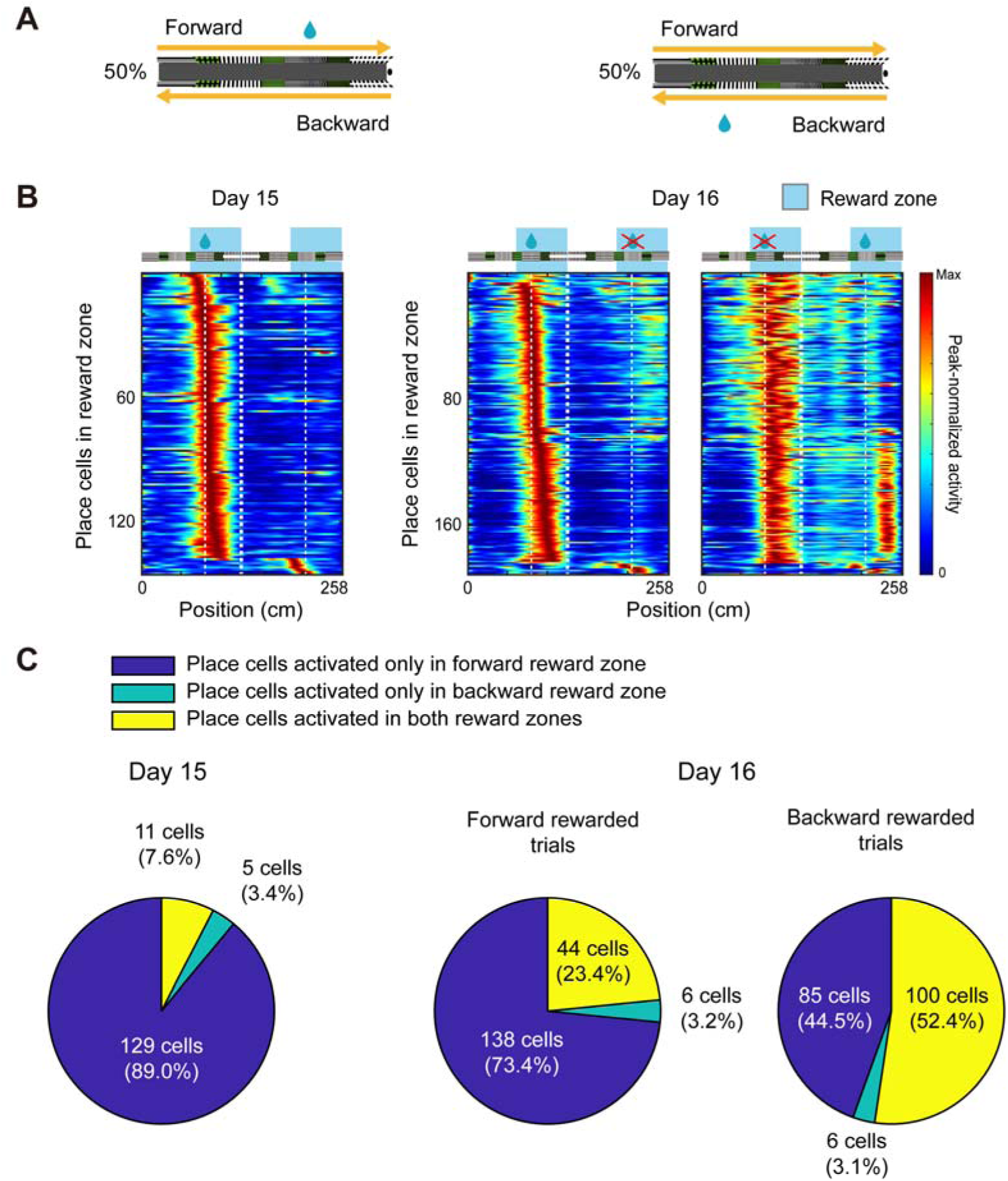
Calcium activity maps in the random reward task. (A) Experimental protocol of the random water reward delivery task. In each trial, mice received water reward from one of two reward locations with a 50% random probability. (B) Left: The activity maps of RSCd-projecting ACC neuronss on day 15 (100% probability, one reward location). Middle: The activity map of RSCd-projecting neurons in the forward reward delivered trials. Right: The activity maps in the backward reward delivered trials. The same set of neurons are shown in the middle and right panels, sorted by the peak activity location in the forward reward task. (C) A pie chart depicting reward place cells proportions.

**Supplementary Figure 5.**
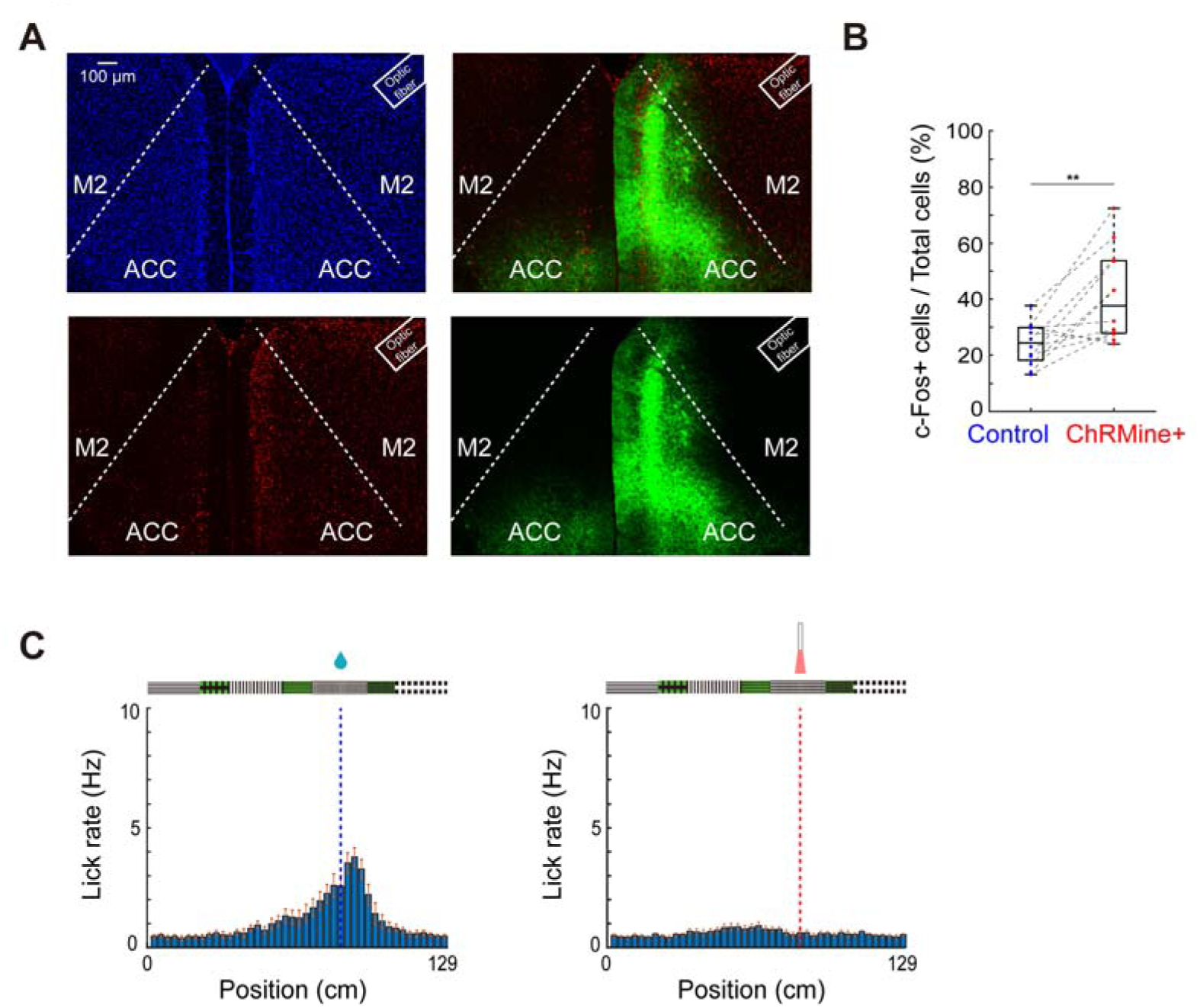
c-Fos staining and lick behavior of mice in the RSCd-projecting neuron activation experiment. (A) Example confocal images of optogenetic activation of RSCd-projecting ACC neurons in a single hemisphere and anti-c-Fos staining. (B) The proportion of c-Fos positive cells in ACC in the optogenetically activated and non-stimulated hemispheres. (C) Lick rate of mice expressing ChRmine in RSCd-projecting neurons during the reward delivery or light stimulation.

**Supplementary Figure 6.**
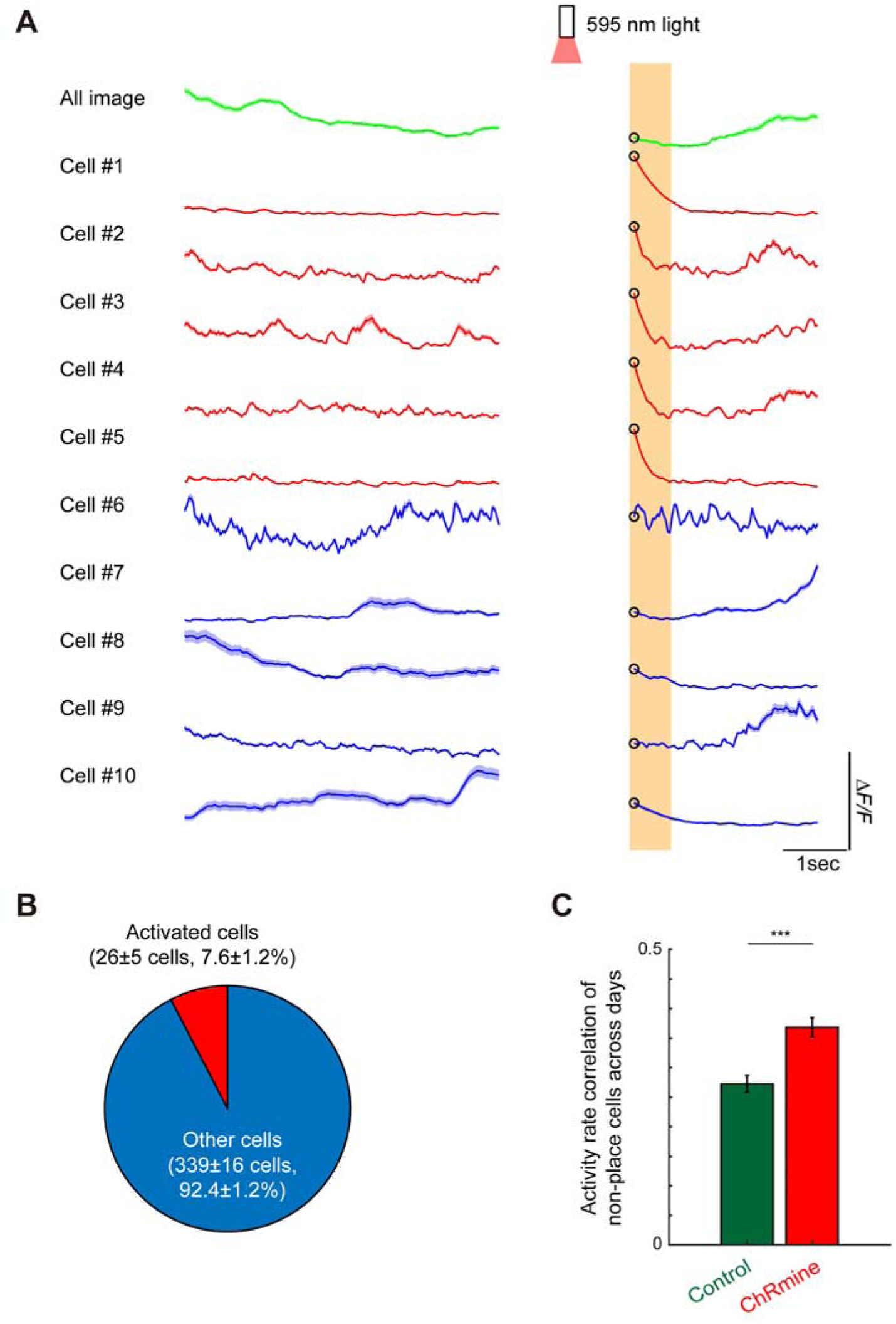
Effects of activation of RSCd-projecting ACC neuron on dorsal hippocampal neuronal activity. (A) Example Ca^2+^ transients of the entire field of view (green), light responsive hippocampal neurons (red) and unresponsive neurons (blue). Shaded areas indicate mean ± SEM (9 ± 3 trains of light stimulation). Ca^2+^ response could not be recorded during the light stimulation due to optical-bleed through of the stimulation light into the detector. (B) The proportion of light responsive and unresponsive dorsal hippocampal neurons. (C) The activity correlation of non-place cells across days (* p < 0.001).

## METHODS

### Animals

Male C57BL/6JmsSlc mice (SLC), aged 12-48 weeks were used in all experiments. Mice were housed in groups on a 12-hour light cycle, provided food and water ad libitum, and maintained at a temperature of 21-23= and humidity of 40-60%. Experiments were performed in accordance with the care and use of laboratory animals approved by Kyoto University.

### Adeno-associated Virus Vector Package

The plasmids of pAAV-CaMKIIα-GCaMP6f (plasmid #100834), pAAV-Synapsin-miRFP670-3XNLS-WPRE (#159952), pAAV-EF1α-DIO-ChRmine-eYFP-KV2.1-WPRE (#130997), pAAV-hSyn-DIO-hM4D(Gi)-mCherry (#44362), pENN-AAV-CamKIIα-Cre-SV40 (#105558), pAAV-hSyn-EGFP (plasmid #50465), pAAV-CAG-tdTomato (#59462) were obtained from Addgene. Retrograde or serotype 9 AAV vectors were packaged and purified according to the protocols available from Addgene. Briefly, AAV vectors were transfected into HEK293T cells, and AAV vector particles were harvested 5 days later. Then, the AAV vector particles were purified by iodixanol gradient ultracentrifugation, and the titers of AAV vectors were measured by qPCR.

### Viral Injections

Mice were anesthetized with gaseous isoflurane (3% induction, 1.5% maintenance) and head fixed in stereotactic apparatus. The virus vector solution was injected at the rate 1 μl⋅h^−1^ using a glass micropipette attached to a microsyringe (MS-10; Ito Corporation, Japan) through a tube filled with liquid paraffin. Injection rate and volume were controlled by an injection pump (KDS100; KD Scientific, MA, USA). The tip of micropipette was left for 10 minutes post-injection to prevent backflow of the virus. For dorsal hippocampus retrograde tracing, 300 nl of AAV2-retro-CAG-tdTomato (titer: 1.5×10^13^ viral genome (vg)⋅ml^-1^) and was injected into the dorsal hippocampus (anterior–posterior: −2.0 mm; medial–lateral: −2.0 mm; dorsal–ventral: −1.6 mm). For ACC projection neuron tracing, 200 nl of AAV2-retro-hSyn-EGFP (titer: 1.5×10^13^ vg⋅ml^-1^), and 150 nl of AAV2-retro-CAG-tdTomato (titer: 1.5×10^13^ vg⋅ml^-1^) were injected into the left dmSTR (anterior–posterior: +0.5 mm; medial–lateral: +1.5 mm; dorsal–ventral: −2.2 mm) and RSCd (anterior–posterior: −2.5 mm; medial–lateral: +0.5 mm; dorsal–ventral: −0.5mm; angle: 20 degree from lateral to medial in the coronal plane), or dmSTR and RSCg (anterior–posterior: −2.5 mm; medial–lateral: +0.3 mm; dorsal–ventral: −0.8 mm), or RSCd and RSCg respectively. For calcium imaging, 250 nl of AAV2-retro-CaMKIIα-GCaMP6f (titer: 2×10^13^ vg⋅ml^-1^) was injected into the left dmSTR (anterior–posterior: +0.5 mm; medial–lateral: +1.5 mm; dorsal–ventral: −2.2 mm), 200 nl of AAV2-retro-CaMKIIα-GCaMP6f (titer: 2×10^13^ vg⋅ml^-1^) was injected into the left RSCd (anterior–posterior: −2.5 mm; medial–lateral: +0.5 mm; dorsal–ventral: −0.5 mm; angle: 20 degrees), 200 nl of AAV2-retro-CaMKIIα-GCaMP6f (Titer: 2×10^13^ vg⋅ml^-1^) was injected into the left RSCg (anterior–posterior: −2.5 mm; medial–lateral: +0.3 mm; dorsal–ventral: −0.8 mm), or 400 nl of AAV9-CaMKIIα-GCaMP6f (titer: 4×10^12^ vg⋅ml^-1^) was injected into the left dorsal hippocampus (anterior–posterior: −2.0 mm; medial–lateral: +2.0 mm; dorsal–ventral: −1.6 mm). It should be noted that our analysis using two different retrograde AAV vectors underestimates the overlap because we cannot ensure that 100% of the projection neurons were infected with the viral vector.

For optogenetic activation experiments, 200 nl of AAV9-EF1α-DIO-ChRmine-eYFP (titer: 2.78×10^13^ vg⋅ml^-1^) was injected bilaterally into ACC (anterior–posterior: 0.0 mm; medial–lateral: ±0.5 mm; dorsal–ventral: −1.45 mm; angle: 15 degrees from lateral to medial in the coronal plane) and 200 nl of AAV2-retro-CaMKIIα-Cre (titer: 2.2×10^13^ vg⋅ml^-1^) was injected bilaterally into RSCd (anterior–posterior: −2.5 mm; medial–lateral: ±0.5 mm; dorsal–ventral: −0.5 mm; angle: 20 degrees). For chemogenetic inhibition experiments, 200 nl of AAV9-hSyn-DIO-hM4D(Gi)-mCherry (titer: 3.6×10^13^ vg⋅ml^-1^) was injected bilaterally into ACC (anterior–posterior: 0.0 mm; medial–lateral: ±0.5 mm; dorsal–ventral: −1.45 mm; angle: 15 degrees) and 200 nl of AAV2-retro-CaMKIIα-Cre (titer: 2.2×10^13^ vg⋅ml^-1^) was injected bilaterally into RSCd (anterior–posterior: −2.5 mm; medial–lateral: ±0.5 mm; dorsal–ventral: −0.5 mm; angle: 20 degrees from lateral to medial in the coronal plane).

### Implantation of prism and lens

More than 2 weeks after virus injections, a stainless-steel head plate (25 mm long, 4 mm wide, 1 mm thick, with a wide circular opening in the center) was implanted onto the skull as previously described (Sato et al., 2016). The center of the opening was targeted at bregma for ACC imaging, and 1 mm posterior to bregma and 1 mm lateral to the midline in the left hemisphere for dorsal hippocampus imaging. For ACC imaging, a right-angle prism was used. A prism-attached optical window (RPB3-1.5-450/2000; OptoSigma, Japan) was prepared as described previously (Sato et al., 2016) with some modifications. To make an optical window targeted to the left hemisphere, a 2.5 mm diameter circular craniotomy was made in the skull and the dura overlaying the right ACC was removed (Ryan J. Low. et al., 2014). The exposed part of the imaging window was fixed with skin adhesive (DHVM12; DERMABOND, USA) and the exposed skull was covered with dental cement. For hippocampal imaging, a GRIN lens (1.0 mm diameter, ∼4.0 mm length; 130-000143; Inscopix, USA) was implanted above the dorsal hippocampus. A 1-mm diameter circle of dura and overlying cortex (anterior-posterior: −2.0 mm; medial-lateral: +2.0 mm) were removed and the GRIN lens was inserted into the brain. Following this, the exposed part of the GRIN lens was fixed with dental cement. For optic fiber implantation, two 0.5 mm diameter circular craniotomies (anterior–posterior: 0.0 mm; medio–lateral: ±1.5 mm) were made on the skull, after which the 200 µm diameter optic fibers covered by cannula (Thorlab, USA) were inserted into the brain with an angle of 60 degrees from lateral to medial in the coronal plane.

### Brain tissue processing and immunohistochemistry (IHC)

Brains were transcardially fixed with 4% paraformaldehyde and were sliced into 100 μm sections with a vibratome to confirm viral expression in ACC projection neurons. For immunohistochemistry, the brains were sliced at a thickness of 50 μm. Prior to staining, slices were blocked in blocking buffer (0.1 M Tris-HCl, 0.15 M NaCl, 0.5% (w/v) blocking reagent (41116100, Roche)) at room temperature (RT) for 1 h. Slices were then washed with TNB-T (0.1 M Tris-HCl (pH 7.5), 0.15 M NaCl, 0.5% (w/v) blocking reagent, 1% Triton (R) X-100) twice for 2 min each and incubated with a monoclonal mouse anti-CaMKIIα antibody (1:250; 6G9, Abcam), or a polyclonal rabbit anti-GAD67 antibody (1:300; GTX 101881), or a monoclonal rabbit anti-c-Fos antibody (1:1,000; 9F6 #2250, Cell Signaling) in TNB-T overnight at 4 °C. The slices were then washed twice for 2 min each and incubated with a goat anti-mouse IgG antibody-Alexa Fluor488 (1:100 diluted in TNB-T; A-11001, Thermo Fisher Scientific), or a goat anti-rabbit IgG antibody-Alexa Fluor488 (1:100; A-11008, Thermo Fisher Scientific), or a goat anti-rabbit IgG antibody-Alexa Fluor594 (1:200; 11012, Thermo Fisher Scientific) at room temperature for 2 h. Finally, slices were incubated with 10 mg/ml Hoechst 33258 (diluted in phosphate-buffered saline; Calbiochem) at room temperature for 5 min for nuclear counterstaining and mounted on glass slides for confocal microscopic imaging.

### Virtual Reality (VR) System

A VR system with a Styrofoam wheel treadmill for head-fixed mice as described previously (Takamura et al., 2021) was used in all experiments. Briefly, head-fixed mice were placed on the treadmill, and an LCD monitor was placed 30 cm in front of the mice to present VR scenes rendered by OmegaSpace 3.1 (Solidray Co. Ltd., Japan). The movement of the treadmill was measured with a USB optical computer mouse (G400, Logitech, Newark, CA) using custom drivers and LabVIEW software (National Instruments, Austin, TX). The velocity signal was converted to analog control voltages (0-5 V) via a D/A converter and fed to a USB joystick controller (BU0836X, Leo Bodner, Northamptonshire, UK) connected to the OmegaSpace computer, to shift the VR scene based on the mouse’s actual movement. Water rewards (5 µl/reward) were delivered by a microdispensing unit (O’Hara & Co., Ltd., Tokyo, Japan) attached to a water delivery tube positioned directly in front of the mouse’s mouth. Behavioral parameters, such as the mouse’s location in the virtual environment, water reward trigger signals, and rotational speed signals of the treadmill, were recorded at 20-ms intervals using custom software in LabVIEW. During calcium imaging, TTL pulses for each frame capture were used to synchronize the imaging and behavioral data.

We designed two types of VR track each with different wall patterns. One track was a linear track and the other a square track. In the linear track, mice run in the same track in opposite directions. When mice arrived at the end of the track, the VR view automatically rotated 180 degrees, and then mice continue running in the opposite direction. In the square track, when mice arrived at a corner, the view automatically rotated 90 degrees, and mice run to the next corner. Each track had a water delivery point, which was delivered as mice arrived.

### Behavior

VR navigation tasks began at least two weeks post-surgery. Prior to the actual VR task, mice were habituated to the VR setup which contained only a gray floor scene, with water reward given randomly. Each day consisted of two habituation sessions. Experiments began after mice ran more than 4000 cm during 20 min.

### *In vivo* calcium two-photon imaging

Ca^2+^-imaging was performed with a Nikon A1MP microscope (Nikon, Tokyo, Japan) controlled by Nikon NIS-elements software. A 910 nm Ti-sapphire laser (MaiTai DeepSee eHP, Spectra-Physics, Santa Clara, CA) was used to excite GCaMP6f. Images of 512 x 512 pixels were acquired at a rate of 15 frames per second using a resonant galvo scanner mounted on the microscope. ACC was imaged using a 16x, NA 0.8 water immersion objective (N16XLWD-PF, Nikon) producing a field of view of 532 x 532 µm. The hippocampus was imaged through a GRIN lens using a 10x, NA 0.45 dry objective (Plan APO 10x/0.45, Nikon) producing a field of view of 425 x 425 µm.

### Image processing and analyses

Calcium images were processed as previously described (Sato et al., 2017). First, a spatiotemporal median filter (7 x 7 x 3 pixels for ACC imaging, 3 x 3 x 3 pixels for dorsal hippocampus imaging) was used to denoise the imaging data. Next, the data were motion-corrected in two steps using a custom MATLAB macro, the first corrects X-Y rigid shifts to remove all motion and the second uses the Lucas-Kanade method for non-rigid frame warping. Next, ROIs and calcium signals of individual cells were extracted using a modified constrained non-negative matrix factorization (CNMF) algorithm, as described in detail elsewhere (Pnevmatikakis, E. A. et al., 2016). If two ROIs had overlapping areas greater than 80%, the ROIs were considered to be the same. All ROIs whose central location was less than 10 pixels from the FOV edges were discarded.

Calcium intensities higher than the threshold (threshold was defined as events exceeding the 97^th^ percentile plus 2.5 SD; and 97^th^ percentile plus 1.5 SD for dorsal hippocampus) in the rising phase of the fluorescence transient were detected as calcium activity event (maximum activity rate equal to 15 Hz).

### Identification of place cells

We calculated spatial information (*I*) to determine which neurons displayed spatial tuning (Skaggs, W. et al., 1992; Souza, B. C. et al., 2018). The *I* of each cell was calculated as follows:

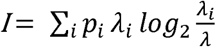

where *λ_i_* is the mean activity in the *i^th^* spatial bin and *p_i_* is the occupancy ratio of the bin, while *λ* is the overall mean activity of the cell. Only neuronal activity during periods of movement were used for place cell related analyses. We defined these periods as the time when the mouse moved at a speed of > 4 cm/s continuously for a duration of > 1 s to reject irrelevant movements, such as grooming and jittering on the treadmill. The entire linear track was divided into 100 bins (bin size = 2.58 cm) and square track was divided into 60 bins (bin size = 1.8 cm). The mean calcium activity rate was calculated for each bin in a session and smoothed by a Gaussian filter (window size = 10 frames, s = 3.5). We defined the smoothed average event frequency as the place activity of each cell during the VR task. Then we calculated *I* and maximum values of place activitys using data that were obtained by randomly rotating the activity during running on each trial. We compared the *I* value and maximum value of place activity with 1000 randomly calculated *I* values and maximum values of place activity for the same cell. If *I* and the maximum value of place activity in real data were >95^th^ percentile of values obtained from randomly permuted data and had an overall mean activity rate more than 0.2 Hz, neurons were considered place cells. If the peak activity of place cells occurred within 7 spatial bins before and after the reward location, place cells were defined as reward place cells.

### Cell alignment across days

To align cells across days, we first averaged the motion-corrected and filter-smoothed calcium imaging data across days. We used the first averaged image as the reference image, and images in subsequent days were aligned to the reference image. Once we obtained the shifted distances between images, the ROIs in different days were shifted according to the shifted distances. If two ROIs on different days overlapped more than 50% and the distance between their centroids was less than 10 pixels, these ROIs were classified as the same cell.

### Logistic regression decoding

We used a logistic regression model to predict reward outcomes based on reward place cell activity in rewarded and non-rewarded trials. The model was constructed as follow:

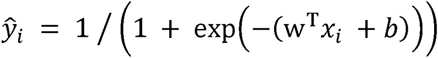

Where *y_i_* is the binary value indicating whether reward was given (*y_i_* = 1) or not (*y_i_* = 0) in *i^th^* trial, *x_i_* is the neural activity vector consisted of peak activity of all reward place cells in *i^th^* trial, w is the weight vector (one weight per neuron) learned by the model, and *b* is the bias (intercept) term. We randomly split all trails into two equal datasets, one of which was used to train the regression model and the other used for decoding. This process was repeated 10 times and an average decoding accuracy score was calculated. Next, to create a shuffled distribution of decoding accuracies to compare against the true value, population activity peaks were randomly shuffled 1000 times.

### Activity rate correlation calculation

Activity rate correlations were calculated as previously described (Leutgeb, J. K. et al., 2005; Tuncdemir, S. N. et al., 2023). Briefly, activity correlations were calculated using a Pearson correlation of place cell activity in all spatial bin (n = 100 bins, bin size = 2.58 cm), between pre and post light stimulation sessions. Only place cells which activated in both sessions were included in the analyses.

### Optogenetic activation

A 595 nm LED light (M595F2; Thorlab) was used to optically stimulate ChRmine-expressing cells. The fiber optic cannula was inserted into the brain and connected to the LED light source via a fiber optic cable (M87L01; Thorlab). The power of the light emitted from the fiber tip was 1-2 mW. A pulse generator (Master 8, A.M.P.I) was used to control the train of LED light pulses, which were delivered at 15 ms/pulse at 20 Hz, with 60 pulses per train.

For c-fos staining experiments, optical fibers were inserted into the ACC of the left hemisphere of mice expressing ChRmine in ACC-RSCd neurons. Two weeks later, mice were stimulated for 10 min with a train of LED light pulses every 30 sec while animals remained in their home cages. Brains were extracted 30 min after stimulation and IHC was performed on the 50 µm brain sections with an anti-c-Fos antibody as described above. The number of c-Fos positive neurons was quantified in the ChRmine expressing area of the left ACC and in the comparable area of the right ACC.

For identifying the light responsive hippocampus neurons, we calculated their calcium activity ± 10 sec from light delivery, and light stimulation period signals were removed. If the neuronal calcium activity reached maximum in 2 sec immediately after light stimulation period and followed a decrease phase, we determine these neurons as light responsive hippocampus neurons.

### Chemogenetic inhibition

To inhibit ACC-RSCd neurons, AAVs were used to express hM4D(Gi) in bilateral ACC-RSCd projection neurons. Five mg/kg of clozapine N-oxide (CNO) (16882; CAYMAN) was administered intraperitoneally at 0.05 mg/ml concentration. VR navigation and calcium imaging began 30 minutes after the injection.

### Statistics

One-tailed or two-tailed Student’s *t*-test was used if the distribution of the group followed a normal distribution. Otherwise, Wilcoxon rank sum tests or Wilcoxon signed-rank test was used. When more than two groups were being compared, analysis of variance (ANOVA) tests were used if the distribution of the groups followed a normal distribution. Otherwise, a nonparametric version of ANOVA (Kruskal-Wallis test) was used. In both parametric and nonparametric ANOVA, p-values were adjusted for post hoc multiple comparisons. Statistical tests were performed in MATLAB. Data are shown as mean ± SEM throughout.

